# FDXR Upregulation by p63/p73 is a Prognostic and Therapeutic Marker of HTLV-1-Associated Adult T Cell Leukemia/Lymphoma

**DOI:** 10.64898/2026.02.25.707207

**Authors:** Lara Gallucci, Maria Josephine, Yasunori Kogure, Alex Ap. Rosini Silva, Vincent Cantaloube-Ferrieu, Marcos Eduardo Souza Abreu, Trusha Ramteke, Phoebe Kibble, Guilherme Lucchesi D Angelo Braz, Rubens de Assis Santos Sebastião, Luiza Amoroso Lambaz, Aijun Lin, Beth Hoare, Felipe Custódio Carvalho dos Santos, Sophie Shackley, Alex Quin, Sofia Luari Moelas Giuzio, Philip A. Lewis, Wujuan Zhang, Youko Nukui, Juliana Pereira, Hebert Fabricio Culler, Takuro Kameda, Kazuya Shimoda, Kate Heesom, Aaron R. Goldman, Ester Cerdeira Sabino, Seishi Ogawa, Fabio Eudes Leal, Mattia Forcato, Keisuke Kataoka, Sheila de Oliveira Garcia Mateos, Iart Luca Shytaj

## Abstract

Infection with human T-lymphotropic virus 1 (HTLV-1) can lead to severe diseases, including adult T-cell leukemia/lymphoma (ATL), for which current therapies are inadequate. HTLV-1 persists chronically through integration of its genome into host DNA and expansion of infected clones via cellular proliferation. Selective elimination of infected cells is the ideal treatment goal but requires identifying cellular markers dysregulated by HTLV-1 that can be exploited therapeutically. Combining metabolomics and proteomics analyses, we identified increased expression of the mitochondrial enzyme ferredoxin reductase (FDXR) as a hallmark of HTLV-1-infected cells. In patient cells, FDXR upregulation mirrored disease severity, was enriched in proliferating tumour cells and predicted worse ATL prognosis. Mechanistically, FDXR upregulation was directly driven by the transactivating isoforms of two p53 family members, TAp63 and TAp73. *In silico* drug susceptibility predictions, followed by targeted cytotoxicity, proliferation and Drug-Seq assays demonstrated that FDXR upregulation is associated with increased sensitivity of ATL cells to repurposed chemotherapeutic agents, particularly fluorouracil. Overall, our findings establish FDXR as a novel biomarker of HTLV-1-associated disease progression and ATL prognosis, elucidate its upstream molecular regulation, and highlight its predictive potential for chemotherapeutic sensitivity of ATL cells.

## Introduction

Adult T Cell Leukemia/Lymphoma (ATL) is an hematological malignancy caused by infection with the human T lymphotropic virus 1 (HTLV-1)^1^. Although ATL is relatively uncommon, occurring in up to 10% of infected individuals after decades of latency, the disease progresses aggressively, particularly in the acute and lymphomatous subtypes, which are characterized by a median survival of less than one year^2^. HTLV-1 infection can also cause other severe conditions, such as HTLV-1–associated myelopathy/tropical spastic paraparesis (HAM/TSP), a chronic and progressive neurodegenerative disorder^3^. Therapeutic options for ATL and HAM/TSP remain inadequate, with the former displaying high relapse rates following chemotherapy, immunotherapy, and even allogeneic stem cell transplant, which is currently the only potentially curative intervention^4^.

Intensive efforts have focused on the identification of molecular biomarkers enabling risk stratification and targeted therapy. These include viral parameters (e.g., HTLV-1 proviral load^5^ and clonality^6^), soluble immune mediators^7^, patterns of somatic alterations^8^ and/or dysregulated expression of host genes^9–11^ and microRNAs^12^. Among these, host genes and pathways which are selectively upregulated in infected cells offer opportunities for targeted therapeutic treatment. A proof of concept is provided by CCR4, which is upregulated in the majority of HTLV-1 infected cells^11^. The monoclonal antibody mogamulizumab, which selectively targets CCR4, has shown remarkable effects in decreasing proviral load and has displayed anti-tumour activity in clinical trials ^13–16^. Nevertheless, overall survival in ATL remains poor, even among responders^15,16^, and no significant clinical benefit could be detected in a recent phase III clinical study of HAM/TSP patients^13^.

Multi-omics integration has become an increasingly powerful strategy for robust biomarker discovery. Studies combining whole-genome sequencing, epigenomic, and transcriptomic data and, more recently, plasma metabolomics, have enabled the identification and validation of key pathways dysregulated in ATL^8,17–20^. On the other hand, proteomic datasets remain underexplored in the field, and the intracellular metabolome of HTLV-1 infected or ATL cells has not been characterized. In this regard, our work on cells infected with the human immunodeficiency virus-1 (HIV-1), which like HTLV-1 is a retrovirus with preferential tropism for activated CD4 T cells, shows that incorporating intracellular metabolomics into multi-omics pipelines can uncover metabolic vulnerabilities amenable to therapeutic targeting^21^.

In the present study we integrate multiple layers of bulk and single-cell omics data from cell models and patient-derived samples and demonstrate that HTLV-1 infected cells upregulate the mitochondrial enzyme ferredoxin reductase (FDXR) as part of a broader metabolic reprogramming toward enhanced mitochondrial activity. *FDXR* mRNA expression correlates with disease progression and poor prognosis in ATL, and is more frequent, at a single cell level, in proliferating tumour cells. Mechanistically, FDXR induction is driven by the infected-cell-specific expression of p53 family members p63 and p73, specifically their trans-activating isoforms (TAp63 and TAp73). By combining gene expression profiling with drug-susceptibility screening, we identify FDXR expression levels as a predictor of sensitivity to several chemotherapeutic agents not previously tested in ATL and include validation of fluorouracil and bleomycin responses *in vitro* and *ex vivo*.

Together, our results uncover FDXR upregulation as a mitochondrial hallmark of HTLV-1 infection and pathogenesis that may serve as a prognostic tool and therapeutic vulnerability to selectively target ATL cells.

## Results

### Upregulation of the mitochondrial enzyme FDXR is a hallmark of HTLV-1 infected cells

To identify cellular markers of metabolic dysregulation associated with HTLV-1 infection, we initially conducted a parallel metabolomic and proteomic analysis using lymphoid cell models (**Supplementary Figure 1A**). We used two HTLV-1–infected cell lines (MT-2 and MT-4), one ATL-derived cell line (SP), and compared them to three uninfected controls (Jurkat, MOLT-4, and CEM-ss). Metabolomic profiling identified multiple mitochondrial pathways as significantly enriched in infected cells, with the electron transport chain, part of oxidative phosphorylation (OXPHOS), emerging as the most significantly enriched pathway in the overall analysis (**Supplementary Figure 1B**). Metabolites represented in the enriched mitochondrial pathways were among the most significantly altered and were predominantly upregulated in infected cells, with the exception of Acetyl−CoA (**Supplementary Figure 1C**). Consistent with these findings, proteomic analysis revealed significant upregulation of pathways involved in fatty acid β-oxidation and mitochondrial Complex I biogenesis (within the OXPHOS system) in HTLV-1 infected cells (**Supplementary Figure 1D, Supplementary Table 1**).

Differential expression analysis of proteomic data, performed filtering the results to proteins with known mitochondrial localization using the MitoCarta repository^22^, uncovered ferredoxin reductase (FDXR) as the most significantly upregulated mitochondrial protein in infected cells (**Figure 1A, Supplementary Table 2**). Functionally, FDXR contributes to lipid homeostasis, iron–sulfur cluster biogenesis, and redox regulation, and depends on flavin adenine dinucleotide (FAD) as a cofactor^23,24^, which our metabolomic analysis revealed as significantly increased in infected cells (**Supplementary Figure 1C**). We thus tested FDXR expression by qPCR, western blotting of isolated mitochondria, and immunofluorescence using a panel of sixteen cell lines as well as activated and resting primary CD4 T cells and PBMC. The results confirmed a significant upregulation of FDXR in HTLV-1–infected and ATL cell lines (**Figure 1B,C**) as well as its mitochondrial localization (**Figure 1C,D**). Two ATL models, MT-1 and ED cells, exhibited elevated *FDXR* mRNA, but not protein levels (**Figure 1B,C**). The increase in FDXR levels was detectable in infected cells irrespective of the expression pattern of viral structural genes and proteins (*Gag/Pol*, p24, gp46) or regulatory transcripts and proteins (*Tax/Rex*, *Hbz*, Tax), as measured by qPCR and western blot (**Supplementary Figure 2A-E**).

**Figure 1.**
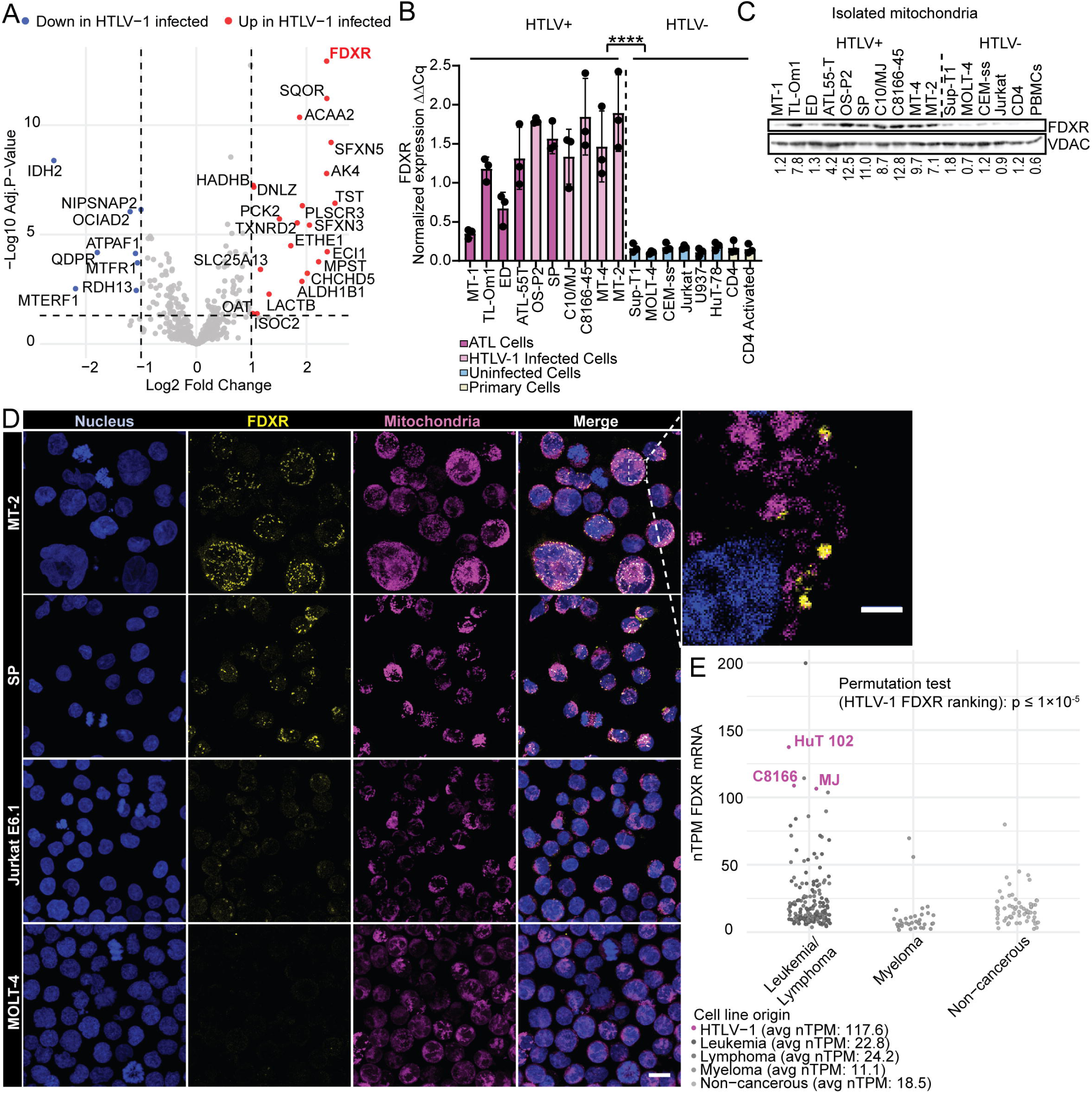
FDXR levels in HTLV-1^+^/ATL, uninfected, and primary blood cell models. **(A)** Three HTLV-1^+^/ATL (MT-2, MT-4, SP) and three uninfected (Jurkat, MOLT-4, CEM-ss) lymphoid cell lines were subjected to proteomic analysis. The volcano plot shows differentially expressed mitochondrial proteins. Differential expression was assessed with the limma package and mitochondrial proteins were filtered using the Mitocarta database v3.0. **(B-D)** FDXR expression in HTLV-1^+^, ATL, and uninfected cell lines and primary cells as measured by qPCR (B), western blot of isolated mitochondria (C), and immunofluorescence (D). In panel B, each dot represents an independent qPCR assay (n = 3), and data are presented as mean ± SD. Relative *FDXR* mRNA levels were calculated by ΔΔCq using *TBP* expression as reference. Data were analyzed by non-parametric Wilcoxon rank-sum test. Panel C shows a representative blot, with relative protein levels reported below each lane and quantified using VDAC as the loading control. Panel D shows representative z-projection images of nuclei (DAPI), FDXR (yellow), and mitochondria stained with MitoTracker (magenta). The boxed region displays a single optical section highlighting mitochondrial localization of FDXR. Scale bars: 10 μm (main) and 2 μm (inset). **(E)** *FDXR* mRNA expression (nTPM) across 259 blood cancer and non-cancerous cell lines, with HTLV-1⁺ cell lines highlighted in magenta. Median ± IQR expression values by cell origin are shown below. The significance of the enrichment of HTLV-1 cell lines among the top cell models for *FDXR* expression in the dataset was assessed by permutation testing (100,000 permutations). RNA-Seq data for all cell lines were retrieved from Protein Atlas. **** p <0.0001.

To further broaden the comparison, we analyzed RNA-Seq data from 259 blood cancer cell lines available in the Human Protein Atlas. The analysis confirmed *FDXR* expression as upregulated in HTLV-1 infected cells (**Figure 1E**). In particular, the HTLV-1–infected cell lines (Hut-102, MJ, and C8166) in the dataset ranked among the top five overall for *FDXR* expression (permutation test p ≤ 1 × 10⁻^5^) (**Figure 1E**). Overall, these data highlight FDXR upregulation as a mitochondrial marker of multiple models of HTLV-1 infection and ATL.

### FDXR expression levels *ex vivo* correlate with disease severity and ATL mortality

To evaluate the physiological relevance of FDXR upregulation, we analyzed transcriptomic data of CD4 T cells and tumour samples from multiple cohorts, including Brazilian and Japanese patients, and complemented these findings with qPCR validation in an independent cohort.

We first conducted a meta-analysis of microarray datasets from four studies^12,25–27^, each including samples from people living with HTLV-1 (PLHTLV), stratified by disease status, alongside healthy controls. In total, the analysis comprised 66 samples from PLHTLV (26 Acute ATL, 20 Chronic ATL, 4 Smoldering ATL, 12 HAM/TSP, and 8 Asymptomatic carriers) and 33 samples from healthy donors. Across all groups of PLHTLV, *FDXR* expression levels were elevated compared with healthy controls, reaching statistical significance in subgroup comparisons for Acute ATL, HAM/TSP, Chronic ATL, and asymptomatic carriers (**Figure 2A**). Notably, *FDXR* expression mirrored disease severity, with Acute ATL and HAM/TSP showing the highest median *FDXR* expression levels (**Figure 2A**).

**Figure 2.**
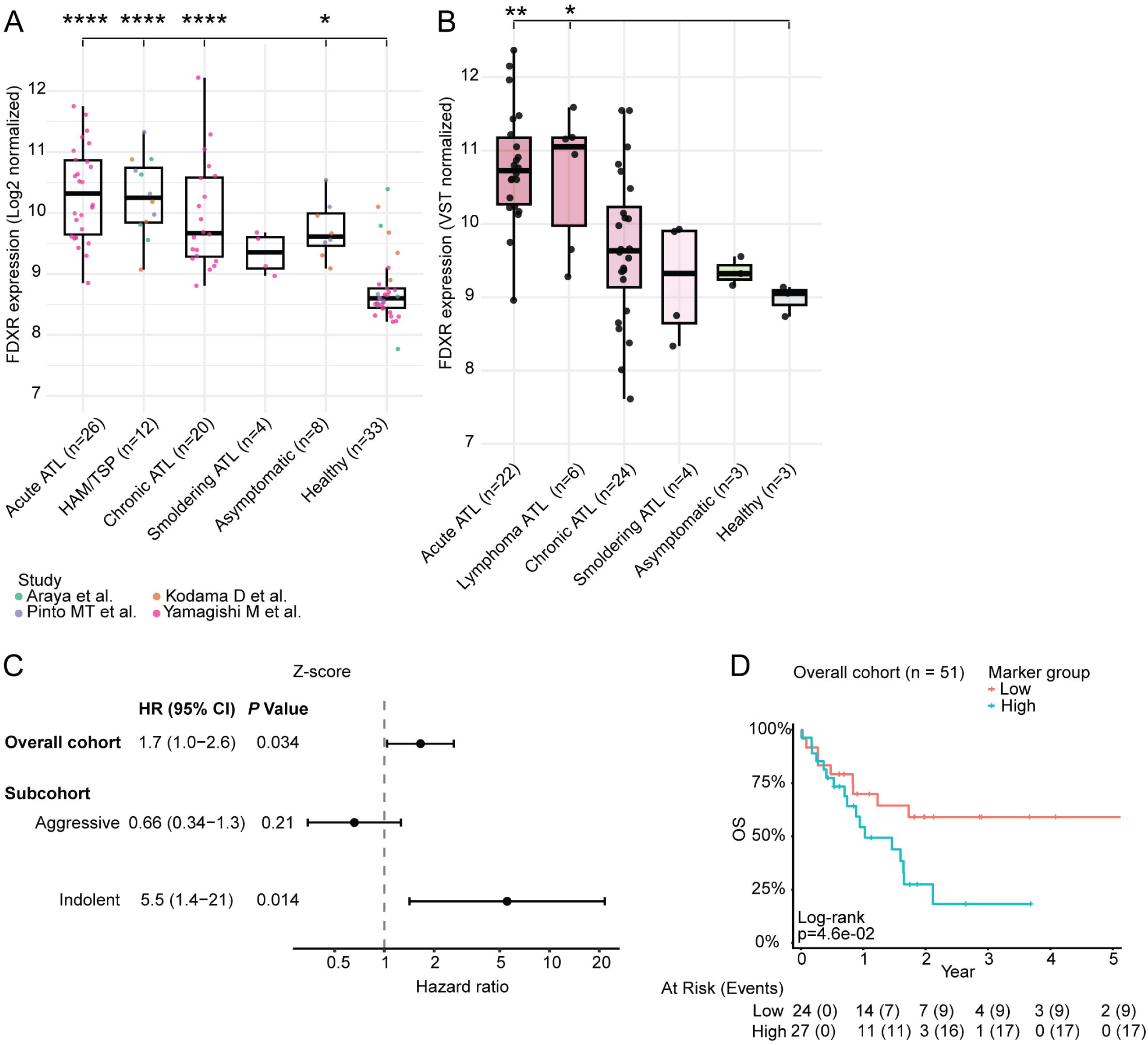
FDXR expression as a predictor of HTLV-1-associated disease severity and ATL mortality. **(A)** Meta-analysis of microarray datasets of CD4 T cells isolated from PLHTLV stratified by disease condition and healthy controls (GSE33615^12^, GSE38537^25^, GSE132666^26^, GSE233437^27^). Data were background-corrected, normalised, and batch-adjusted using the limma^54^ and sva^56^ (ComBat) packages. Each dot depicts Log2 normalized *FDXR* expression levels from a single donor, while black lines indicate median expression levels of each subgroup. Differences in *FDXR* expression were tested using a linear mixed-effects model, with disease status as a fixed effect and study as a random intercept. Post hoc contrasts comparing each disease group with uninfected controls were performed using study-adjusted estimated marginal means (using the emmeans package), followed by Holm correction for multiple testing. **(B)** RNA-Seq data (EGAD00001001411^8^) from tumour and CD4 T cells isolated from PLHTLV stratified by disease condition and healthy controls. Each dot depicts variance-stabilized (VST) FDXR expression levels from a single donor, while black lines indicate median expression within each subgroup. Differential expression between each subgroup and healthy controls was tested using DESeq2. **(C–D)** Association of FDXR expression with overall survival in ATL patients. (C) Cox proportional hazards analysis of continuous Z-score–normalized *FDXR* expression levels in the full cohort and in aggressive (acute, lymphoma) and indolent (chronic, smoldering) ATL subtypes. (D) Kaplan–Meier analysis of overall survival in patients stratified into high and low *FDXR* expression groups based on the median Z-score. *FDXR* expression values were derived from RNA-seq dataset EGAD00001001411 and analysed in ATL patients with available survival data (n = 51^8,66^). Overall survival in (C–D) was censored at hematopoietic stem cell transplantation, and survival differences in (D) were assessed using the log-rank test. **** p <0.0001; ** p <0.01; *p <0.05.

A similar pattern was observed upon reanalysis of RNA-Seq data derived from a dataset^8^ of 59 samples from PLHTLV (22 Acute ATL, 6 Lymphoma ATL, 24 Chronic ATL, 4 Smoldering ATL, 3 Asymptomatic carriers) and 3 healthy controls. Also in this case, median *FDXR* expression was higher in HTLV-1^+^ samples and increased with disease severity, showing statistically significant differences from healthy controls in Acute and Lymphoma ATL samples (**Figure 2B**). Consistent with these findings, qPCR analysis of *FDXR* expression in isolated CD4 T cells from our cohort of blood donors in São Caetano do Sul, Brazil (**Supplementary Table 3, Supplementary Figure 3A,B**), confirmed significantly higher FDXR expression in PLHTLV compared with healthy donors (**Supplementary Figure 3B**). Although median FDXR levels also increased with disease severity in this cohort (**Supplementary Figure 3C**), differences between clinical subgroups did not reach statistical significance, likely reflecting the smaller number of individuals with advanced disease (one acute ATL and nine HAM/TSP patients), compared with the other cohorts analyzed.

We next assessed the prognostic significance of FDXR expression levels on ATL patient survival by integrating RNA-Seq data (**Figure 2B**) with clinical outcome information available from the same cohort (n=51). A Cox model on continuous, z-score–normalized, *FDXR* expression revealed a significant link between higher *FDXR* levels and reduced survival (HR = 1.7, P = 0.034; **Figure 2C**). Incorporating ATL subtype stratification (aggressive [acute/lymphoma] and indolent [chronic/smoldering] subtypes) further showed that *FDXR* expression was significantly associated with worse prognosis within indolent subtypes of ATL (HR = 5.5, P = 0.014; **Figure 2C**). This likely reflects the broader dynamic range of FDXR expression in indolent ATL (**Figure 2A,B**), that can enable greater prognostic discrimination than in aggressive disease, where *FDXR* upregulation and clinical outcomes are more uniform. Finally, Kaplan–Meier survival analyses based on two-and four-group stratifications of *FDXR* expression also demonstrated poorer outcomes among patients with higher *FDXR* levels, reaching statistical significance in the two-group comparison (**Figure 2D**; **Supplementary Figure 3D**).

Taken together, these data show that FDXR expression levels in PLHTLV correlate with disease progression and worse ATL prognosis.

### FDXR upregulation is driven by ectopic expression of TAp63 and TAp73

HTLV-1 infection is known to dysregulate the activity and expression of multiple host genes, including transcription factors (TFs)^28^. To identify putative host mediators of *FDXR* upregulation we first conducted an *in silico* analysis of differential TF activity across leukemia and lymphoma cell lines, including HTLV-1–infected models, using RNA-Seq data retrieved from the Human Protein Atlas. Cells were stratified in two groups according to *FDXR* expression levels (**Figure 3A**). TF activity was inferred from downstream target gene expression using the VIPER algorithm^29^ in combination with a regulatory network of TF–target interactions derived from curated databases (DoRothEA^30^) and interactions inferred directly from the dataset^31^ (**Figure 3A**). The results highlighted the TF activity of two members of the p53 family, p63 and p73, as significantly upregulated in cells expressing higher FDXR levels (**Figure 3B**). Of note, p63 activity was the second most upregulated, ranking after PRDM1, which encodes for a transcriptional repressor of p53^32^. This pattern suggests a potential shift away from p53 dependence toward increased reliance on the p63 and p73 members of the family.

**Figure 3.**
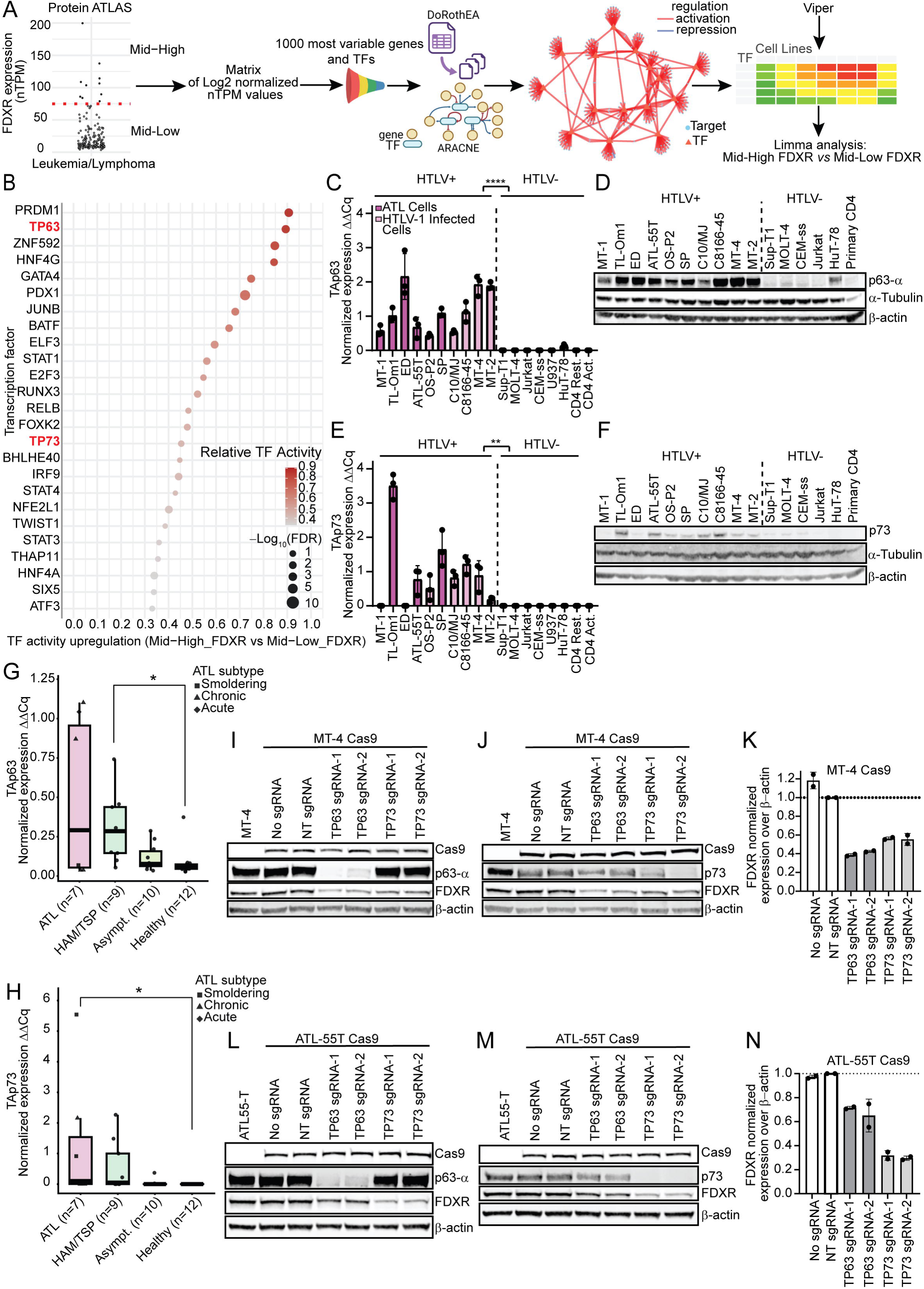
Regulation of FDXR by TAp63 and TAp73 in HTLV1^+^/ATL cells. **(A,B)** RNA-Seq data retrieved from Protein Atlas were stratified by FDXR expression (≥75 nTPM = Mid-high_FDXR, <75 nTPM = Mid-low_FDXR). Differential transcription factor (TF) activity between groups was inferred using VIPER^29^, based on regulatory networks constructed with DoRothEA^30^ and ARACNe. The bubble plot in (B) ranks TFs based on their differential activity in the two groups. The colour of the dots reflects relative TF activity, while the size of the dots represent -Log10 adjusted p values (FDR) calculated with the limma package. **(C–F)** Expression of *TAp63* mRNA (C), p63α protein (D), *TAp73* mRNA (E) and p73 protein (F) in HTLV-1^+^, ATL, and uninfected cell lines and primary cells. Relative mRNA expression was quantified by qPCR; protein expression was assessed by western blot. In panels (C,E), each dot represents an independent qPCR assay (n = 3), and data are presented as mean ± SD. Relative mRNA levels were calculated by ΔΔCq using *TBP* expression as reference. Data were analyzed by non-parametric Wilcoxon rank-sum test. **(G,H)** Relative expression of TAp63 (G) and TAp73 (H) mRNA in primary CD4 T cells from PLHTLV stratified by disease condition and healthy controls. Relative mRNA expression was quantified by qPCR using the ΔΔCq method and *TBP* expression as reference. Each dot represents a donor and data are depicted as median ± IQR. Data were analyzed by non-parametric Kruskal–Wallis test followed by Dunn’s post hoc test with Benjamini–Hochberg correction for multiple comparisons. **(I,N)** Effect of TP63 and TP73 knockout on FDXR expression. The HTLV-1–infected MT-4 cell line (I-K) and the ATL-derived ATL-55T cell line (L-N) were stably transfected with Cas9 and subsequently transduced with lentiviruses expressing either a non-targeting (NT) sgRNA or sgRNAs targeting TP63 or TP73. Stable knockout cell lines were lysed and used to assess protein expression by Western blot. Panels I, L show p63 and FDXR expression in MT-4 and ATL-55T cells, respectively. Panels J, M show p73 and FDXR expression in MT-4 and ATL-55T cells, respectively. Wild-type and non-transduced Cas9-expressing cells served as additional controls. Panels K, N show relative FDXR protein levels normalized using actin as a loading control and the NT sgRNA condition as reference level. Barplots in K, N show mean ± SD for each condition. N = 2. **** p <0.0001; ** p <0.01; *p <0.05.

The *FDXR* promoter harbours a responsive element that can be bound by all members of the p53 family^33^, potentially providing a direct mechanistic link with *FDXR* upregulation. We thus examined their expression in HTLV-1^+^/ ATL cell lines, as well as in primary CD4 T-cells isolated from PLHTLV, using uninfected cell lines and primary CD4 T cells as controls. Because only the TA isoforms of p63 and p73, which retain an intact N-terminal transactivation domain, have been reported to bind the FDXR promoter in colorectal cancer cells^33,34^, we quantified isoform-specific expression using qPCR assays designed to distinguish TA from ΔN variants (**Supplementary Figure 4A**). *TAp63* mRNA and p63 protein were consistently expressed in all HTLV-1^+^/ATL cell models, whereas uninfected cells largely lacked *TAp63*, with only modest expression detected in Hut78 cells (**Figure 3C,D**). Similarly, *TAp73* mRNA and p73 protein were upregulated in HTLV-1^+^/ATL cells, consistent with previous reports^19^, with the exception of MT-1 and ED cells (**Figure 3E,F**), which are also characterized by low or absent FDXR upregulation (**Figure 1B,C**). ΔNp63 isoforms were undetectable across all models (**Supplementary Figure 4B**), while ΔNp73 expression mirrored that of TAp73 (**Supplementary Figure 4C**). In contrast, *p53* mRNA and protein levels showed no significant differences between infected and uninfected cells (**Supplementary Figure 4D,E)**, in agreement with our TF activity analysis. Consistent with these findings, RNA-Seq data^8^ revealed significantly increased expression of *p63* and *p73* in Acute, Lymphoma and Chronic ATL patients (**Supplementary Figure 4F,G**). Moreover, qPCR analysis of primary CD4 T cells of PLHTLV showed *TAp63* and *TAp73* as significantly increased in HAM/TSP and ATL patients, respectively (**Figure 3G,H**).

To further elucidate the mechanistic link between TA isoform expression and FDXR transcription, we performed chromatin immunoprecipitation (ChIP) followed by qPCR using primers specific for the p53-responsive element within the FDXR promoter^33^. In line with expression patterns, TAp63 showed specific binding to the FDXR promoter in HTLV-1^+^/ATL cells, whereas ΔNp63 showed minimal or no enrichment above IgG controls (**Supplementary Figure 4H**).

Building on previous reports that TP73 expression in ATL cells can be induced by the IRF4/BATF3^19,35^ complex, we investigated whether TP63 can also serve as a target for this complex. To this end, we extended our *in silico* analysis of TF activity (**Figure 3A**) to identify upstream regulators of TP63 and TP73. Strikingly, IRF4 emerged as one of the strongest predicted upstream activators of both genes (**Supplementary Figure 5A**). Moreover, analysis of ChIP-seq data generated in ATL cells^35^ revealed binding peaks for both BATF3 and IRF4 at the TP63 locus (**Supplementary Figure 5B**). Scanning the sequence of the TP63 locus revealed a canonical AP1–IRF composite (AICE) motif (**Supplementary Figure 5C**), belonging to a class of regulatory elements known to mediate cooperative binding of IRF4 and BATF3^36^. The AICE motif lay within 50 bp of IRF4 and BATF3 binding peaks (**Supplementary Figure 5C**), overlapped with an active enhancer (FANTOM5 collection^36,37^) and was marked by enrichment of H3K27ac and BRD4 binding. Overall, this was consistent with open chromatin and active enhancer activity (**Supplementary Figure 5B**) suggesting that the IRF4/BATF3 complex can bind both TP63 and TP73 and provide a joint upstream regulation of both genes.

Finally, to provide a direct mechanistic link between FDXR upregulation in HTLV-1^+^/ATL cells and p63/p73, we generated stable CRISPR–Cas9 knockouts in MT-4 (**Figure 3I-K**) and ATL-55T (**Figure 3L-N**) cells. In both models, loss of either p63 or p73 resulted in markedly reduced FDXR protein levels (**Figure 3I-N**).

Taken together, these data demonstrate that FDXR upregulation in HTLV-1^+^/ATL cells is driven by selective expression of TAp63 and TAp73.

### *FDXR* transcription is enriched in proliferating cells of ATL patients

In addition to its role in maintaining mitochondrial homeostasis, FDXR can act as a regulator of cell proliferation^38^. We thus hypothesized that in ATL, increased FDXR expression may preferentially mark proliferating tumour cells. We examined the single-cell expression pattern of *FDXR* in primary cells, leveraging a previously published^39^ scRNA-seq dataset generated using PBMCs of healthy donors and PLHTLV, including ATL patients. After dimensionality reduction and clustering (**Figure 4A**), we focused our analysis on ten clusters of T/NK cells (**Supplementary Figure 6**) from individuals with chronic ATL (samples ATL-1, -2, -3, -6, and -7) and lymphoma (sample ATL-4). These samples showed the highest average frequency of cells with active FDXR gene expression (**Figure 4B**), consistent with our observation that FDXR upregulation increases with disease severity (**Figure 2A,B**). Single-cell analysis further revealed enrichment of active *FDXR* gene expression within the proliferating cell cluster (cluster 8) in all but one ATL sample (**Figure 4C**). The proliferative identity of this cluster was supported by its overlap with cells in the S and G2/M phases of the cell cycle (**Figure 4A**) and by the expression of canonical proliferation-associated marker genes (**Supplementary Figure 7**). Moreover, the relative expression of *FDXR* was more enriched in proliferating cluster 8 as compared to that of CADM1 and Hbz (**Figure 4D**), which are established markers of HTLV-1 infection and ATL^9,39^.

**Figure 4.**
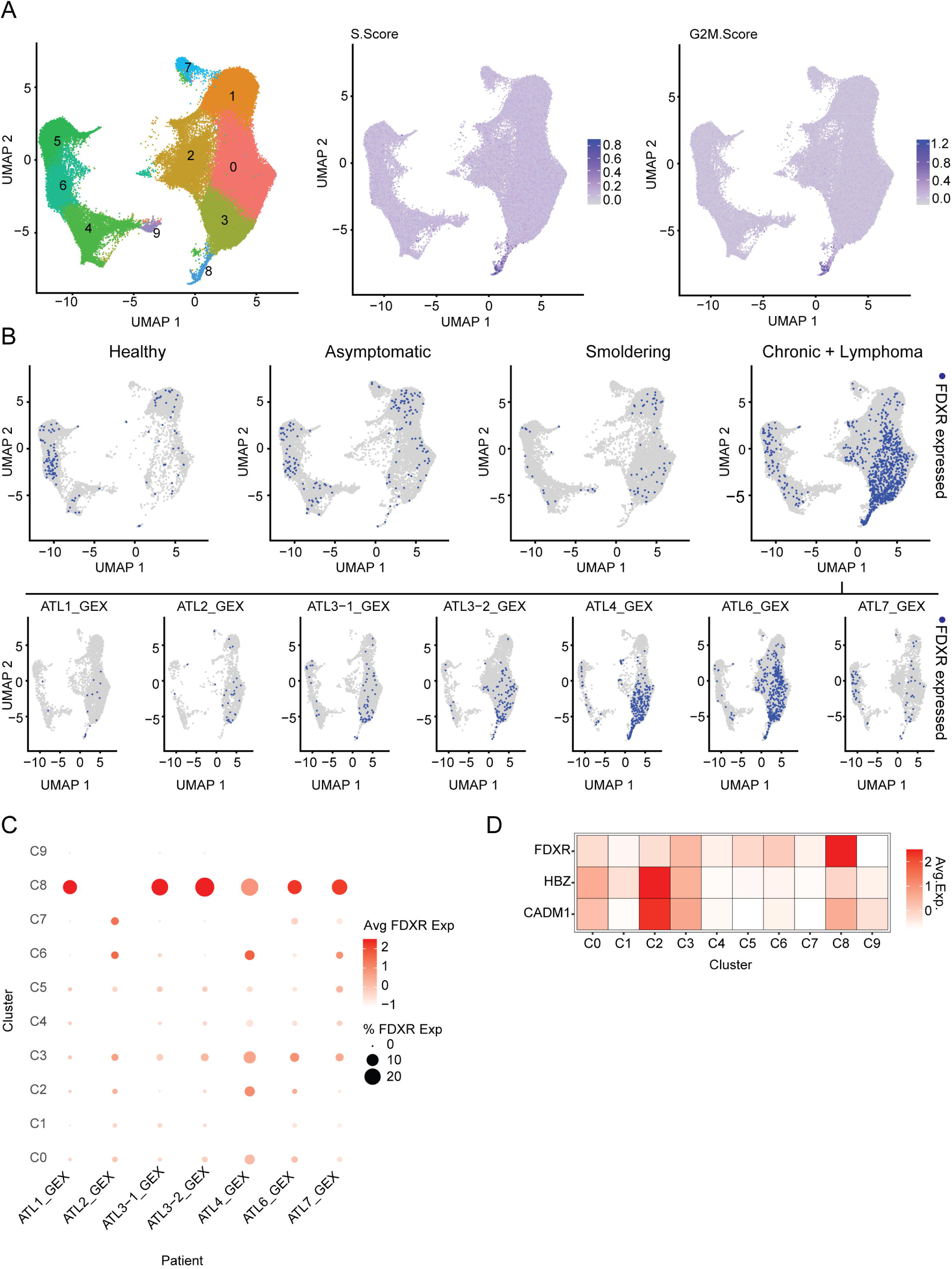
Single-cell *FDXR* expression pattern in blood cells of PLHTLV. Raw scRNA-seq data were retrieved from study PRJEB47382^39^, comprising 17 samples from 16 donors (3 healthy donors, 4 asymptomatic HTLV-1 carriers, 3 smoldering ATL, 5 chronic ATL, and 1 lymphoma-type ATL). **(A)** UMAP visualization of 10 T/NK cell clusters (C0-C9) identified across the dataset (91,436 cells total). Proliferating cells are highlighted by S-phase and G2/M-phase cell-cycle scores, computed with Seurat. **(B)** Distribution of FDXR-expressing cells across sample categories: healthy donors (average FDXR^+^ = 1.20%), asymptomatic carriers (average FDXR^+^ = 0.75%), smoldering ATL (average FDXR^+^ = 0.47%), and chronic/lymphoma ATL (average FDXR^+^ = 2.29%). The UMAP visualizations of individual ATL samples (ATL1–ATL7) display the localization of FDXR-expressing cells within the T/NK subset of each sample. **(C)** Dot-plot showing the frequency and relative expression levels of *FDXR* across clusters (C0–C9) in each ATL sample. **(D)** Heatmap comparing in ATL samples the average expression of FDXR and of the known infection markers^9,39^ HBZ and CADM1. The heatmap shows scaled average expression levels of each gene across clusters (C0–C9). The expression of each gene within each cluster was quantified by aligning reads to a custom reference transcriptome generated by merging the human GRCh38 genome with the HTLV-1 genome (GenBank: AB513134.1). Cluster 8 denotes proliferating cells (Supplementary Figure 7).

Overall, this analysis shows that FDXR expression is selectively enriched at the single-cell level in proliferating cells from ATL patients.

### FDXR expression levels predict sensitivity to selected chemotherapeutics in vitro and *ex vivo*

Baseline FDXR expression and its upregulation upon treatment has been linked to increased sensitivity to ROS-generating chemotherapeutics in colorectal cancer^33,40^. To broadly assess which chemotherapeutics could leverage increased FDXR expression, we analysed cytotoxicity and gene-expression profiles from the Cancer Therapeutics Response Portal (CTRP)^41^. The analysis of the entire dataset (481 compounds tested in up to 835 cell lines) revealed that FDXR expression was significantly associated with enhanced sensitivity to five agents (**Figure 5A**). Restricting the analysis to hematopoietic cell lines identified fluorouracil and bleomycin A2 as the compounds with the lowest dose-response area under curve (AUC) values, a pattern also observed in the only HTLV-1 infected cell line (MJ cells) in the dataset (**Figure 5B)**.

**Figure 5.**
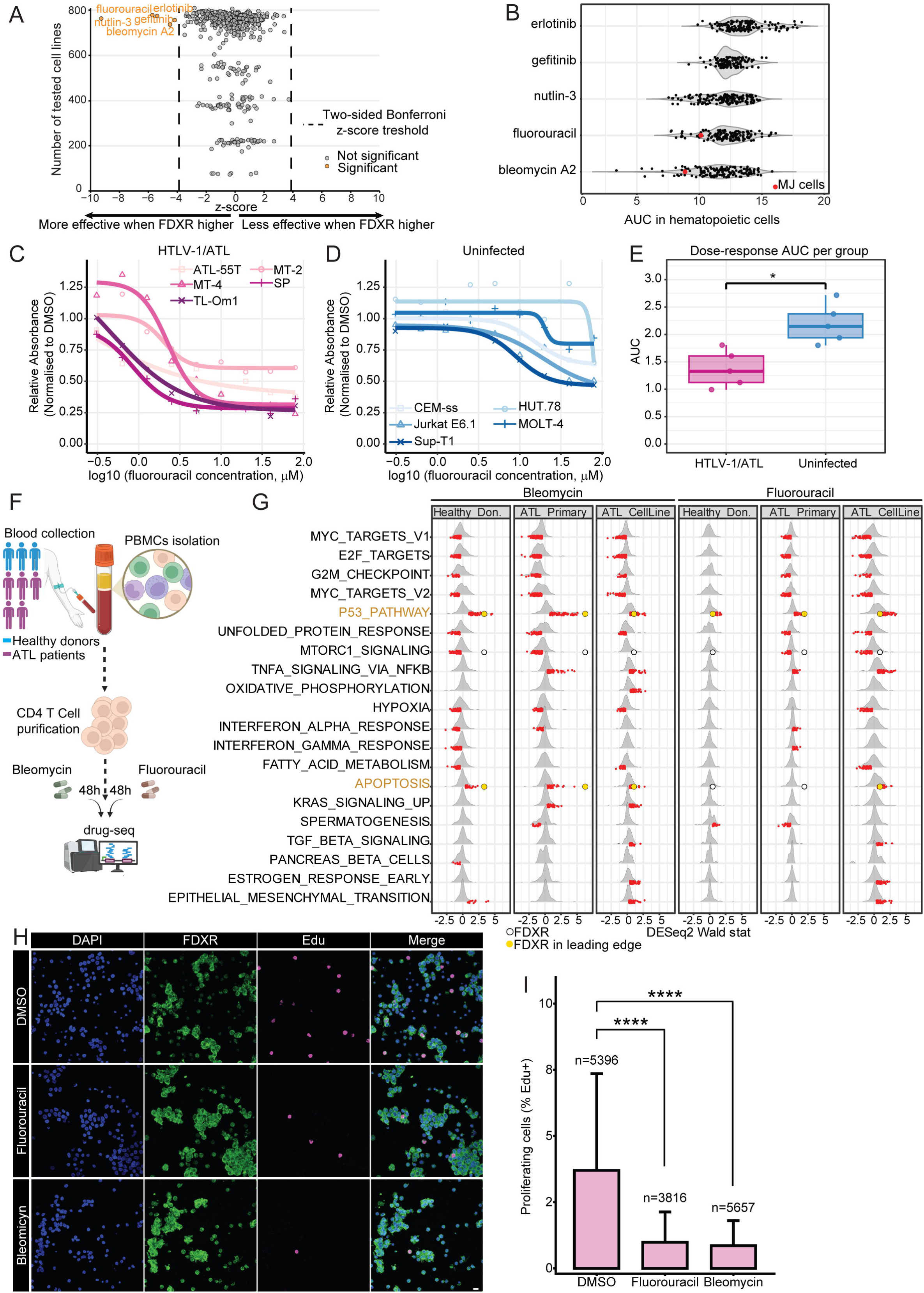
Chemotherapeutics leveraging FDXR expression in HTLV-1^+^/ATL cells. **(A,B)** Data from the Cancer Target Discovery and Development (CTD²) Network^41,73^ were used to assess associations between *FDXR* mRNA expression and sensitivity to 481 compounds across cancer cell lines (up to 835 cell lines per compound). In panel A, the scatter plot shows the association between *FDXR* expression and drug sensitivity for each compound, expressed as a Pearson correlation z-score (x-axis) versus the number of cell lines tested (y-axis). Significance thresholds were defined using a Bonferroni-corrected two-tailed z-test. Panel B shows the distribution of area under the curve (AUC) values in hematological and lymphoid cell lines for compounds significantly correlated with *FDXR* expression. Lower AUC values indicate increased drug sensitivity. Each dot represents a cell line, with the HTLV-1 infected MJ cell line highlighted in red. **(C-E)** HTLV-1^+^/ATL and uninfected cell lines were treated with serial dilutions of fluorouracil, and cell viability/proliferation was assessed by MTS assay 72 h post-treatment. Absorbance values were normalized to DMSO controls, and dose–response curves were fitted for each cell line using four-parameter logistic nonlinear regression (C-D). The fitted curves were used to calculate the AUCs depicted in panel E. In panel E, each dot represents a cell line and data are depicted as median ± IQR. Group-wise AUC comparisons were conducted using an unpaired Wilcoxon rank-sum test. (**F-G**) Primary CD4 T cells isolated from five ATL patients (1 acute, 3 chronic, 1 smoldering) and three healthy donors, as well as two ATL cell lines (SP and TL-Om1), were treated with fluorouracil or bleomycin for 48 h. DMSO-treated samples served as parallel controls. RNA was extracted and subjected to Drug-Seq analysis (F). Ridge plots in panel G show the distribution of DESeq2 Wald statistics for genes within the top 20 Hallmark pathways (top 10 per drug by maximum |NES|) as calculated by GSEA (Supplementary Figure 8D). Red ticks indicate leading-edge genes. FDXR is highlighted in gold when present in the leading edge or indicated by an open circle when present in a pathway but not in the leading edge. **(H-I)** Primary CD4 T cells isolated from three chronic ATL patients were treated with fluorouracil or bleomycin (10μM for each drug) for 72 h. Cells were fixed with 4% PFA and stained with the Click-iT Plus EdU kit to detect proliferation as well as with an anti-FDXR antibody. Panel H displays representative z-projection images showing nuclei (DAPI, blue), FDXR (green), and proliferating cells identified by EdU incorporation (magenta). Scale bar: 10 μm. The barplot in panel I shows the percentage of proliferating (EdU-positive) cells. The total number of cells analyzed per condition is indicated above each bar. Data were analyzed using a binomial generalized linear mixed-effects model with donor as a random effect and Dunnett-adjusted comparisons to DMSO. *p <0.05; ****p <0.0001.

Both fluorouracil and bleomycin A2 are approved chemotherapeutic agents, however, neither has been evaluated in ATL to date. We therefore assessed their effects on cell viability and proliferation by MTS assay in HTLV-1^+^/ATL cell models, using uninfected lymphoid cells as controls. Fluorouracil demonstrated increased efficacy in HTLV-1^+^/ATL cell lines, as indicated by significantly lower dose-response AUC values (**Figure 5C-E**). Treatment with bleomycin was also associated with lower median dose-response AUC values in HTLV-1^+^/ATL cell lines, although this difference was not statistically significant (**Supplementary Figure 8A-C**).

We next evaluated the effects of fluorouracil and bleomycin in primary ATL cells, by performing Drug-Seq on CD4 T cells from five ATL donors (one acute, three chronic, and one smoldering) and three healthy donors (**Figure 5F**). Two ATL cell lines (SP and TL-Om1) were also assayed as controls. Each drug was tested at two concentrations (5 and 10μM). Gene set enrichment analysis using Hallmark pathways revealed that bleomycin significantly perturbed a greater number of pathways as compared to fluorouracil (**Figure 5G, Supplementary Figure 8D**). However, fluorouracil treatment in primary ATL cells and ATL cell lines affected a larger number of pathways as compared with cells of healthy donors (**Figure 5G, Supplementary Figure 8D**), in line with its higher selectivity in dose-response MTS experiments (**Figure 5C-E**). Of note, both bleomycin and fluorouracil significantly affected pathways associated with cell proliferation and cell death, particularly the p53 pathway (**Figure 5G, Supplementary Figure 8D)**. Moreover, FDXR expression was induced by both treatments and FDXR emerged as a leading-edge gene in two pathways following bleomycin or fluorouracil treatment (P53 pathway and APOPTOSIS) (**Figure 5G**). Although in primary cells the pathway-level effects of bleomycin were less selective than those of fluorouracil (**Figure 5G, Supplementary Figure 8D**), at single gene level, FDXR was the most significant differentially expressed gene (DEG) in primary ATL cells treated with each bleomycin concentration (**Supplementary Figure 8E,F)**.

Finally, to validate the antiproliferative effects associated with FDXR upregulation following exposure to fluorouracil and bleomycin, we combined immunofluorescence staining for FDXR with Click-EdU incorporation assays in primary CD4 T cells from ATL donors. The results showed a significant reduction in the frequency of proliferating cells upon treatment with either fluorouracil or bleomycin as compared to DMSO controls (**Figure 5H,I**).

Overall, these data indicate that repurposed chemotherapeutics leveraging FDXR expression can inhibit proliferation of ATL cells *in vitro* and *ex vivo*.

## Discussion

Our findings identify FDXR upregulation as a novel marker of HTLV-1 infection, as well as disease severity and prognosis, thus extending its role beyond solid cancers. In colorectal cancer, FDXR has been linked to p53-dependent pro-apoptotic activity^40^ and proposed as a biomarker for predicting treatment outcome^42^. A similar approach may be applicable to ATL, provided the development of a standardized assay to define FDXR upregulation thresholds. In terms of disease outcome, the prognostic value of FDXR upregulation in our retrospective ATL cohort was significant only in indolent cases (chronic and smoldering). This pattern likely reflects the distribution of FDXR expression across ATL subtypes. While most acute and lymphoma cases exhibit elevated FDXR levels, indolent cases show greater heterogeneity, ranging from marked upregulation to little or no increase relative to healthy donor controls. Since FDXR is preferentially transcribed in proliferating ATL cells, its upregulation in indolent patients may signal atypically high proliferative activity, thereby predicting poorer outcomes. Conversely, in aggressive ATL, where high proliferation is common, further increases in FDXR would offer limited prognostic insight.

The broad availability of fluorouracil and bleomycin, both part of the WHO Essential Medicines List, support their potential for repurposing. However, although in our assays ATL cells displayed increased sensitivity to fluorouracil relative to other hematologic models, their responses remain within the intermediate response range previously reported for colorectal cancer cells^43^, suggesting the need of combination treatments for optimal efficacy. In this regard, targeting anti-apoptotic BCL-2 family proteins is a promising strategy against ATL^44,45^ and may enhance the pro-apoptotic effects associated with FDXR upregulation. More broadly, FDXR upregulation can serve as a basis for targeted drug selection to overcome the well-known chemoresistance of ATL. Notably, nutlin-3, one of the highest-ranking compounds in our predictions, has been previously shown to exert selective toxicity toward wt-p53 ATL cells^46^.

The upregulation of FDXR in HTLV-1–infected cells is driven by p63 and p73, consistent with their ability to bind the FDXR promoter^33^. Whereas enhanced fluorouracil sensitivity in colorectal cancer cells with high FDXR expression was initially reported only in the presence of wt-p53^40^, our data indicate that in ATL cells this can occur even when p53 is absent (*e.g.* in ATL-55T cells). We further show that TAp63 and TAp73 are the isoforms upregulated in infected cells, in agreement with previous reports linking HBZ to TAp73 induction^19^. Of note, TAp63 upregulation by HTLV-1 infection had not been described previously. We find that HTLV-1–infected cells express only the TAp63 isoform, with ΔNp63 completely absent. Given the established role of p63 isoform balance in stem-cell regulation and epidermal biology^47^, it is intriguing to speculate that this dysregulation may be linked to the skin manifestations commonly seen in ATL.

By identifying a novel AICE motif^48^ bound by IRF4/BATF3 in ATL cells and overlapping an enhancer within the p63 locus, we show a potential joint regulation of p63 and p73 by this signaling axis, which had previously been linked only to the latter^35^. IRF4 induction is a recognized hallmark of HTLV-1 infection^8,35^, can be driven by both Tax and HBZ^49^, and has been proposed as a unifying therapeutic target^35,50^. The potential link between IRF4 activation and FDXR upregulation through p63 and p73 would explain our observation that FDXR is upregulated in both ATL and HAM/TSP patients, which typically exhibit high expression of HBZ and Tax, respectively^1,3^. Overall, the pathway here proposed linking IRF-4/p63/p73/FDXR upregulation, cellular proliferation, and elevated mitochondrial metabolism could help contextualize several independent findings in the field, such as increased ROS levels in infected cells^51^, the high proliferative capacity of ATL cells^1,39^, and metabolic dysregulation induced by HTLV-1^19^.

One limitation of our work is that the analyses were mostly confined to CD4 T cells. Although this population harbors the majority of the proviral load, other cell types can also be infected by HTLV-1^1^, and their FDXR expression profiles would warrant investigation in future studies. Moreover, despite the long-standing clinical use of fluorouracil and bleomycin, their potential repurposing for a hematological malignancy such as ATL would necessitate dedicated *in vivo* studies or early-phase exploratory clinical trials. Finally, although FDXR upregulation and its underlying mechanism were supported by data from multiple independent cohorts, the donors from which Drug-Seq samples were obtained included a single acute ATL case. Given that acute and lymphoma ATL subtypes exhibit the highest median FDXR expression, it is plausible that the efficacy of fluorouracil and bleomycin may be higher in aggressive subtypes characterized by elevated FDXR levels.

Taken together, our study reconstructs a molecular pathway converging on FDXR upregulation in HTLV-1 infected cells, revealing its potential as a prognostic marker and as a candidate determinant for chemotherapy selection in ATL.

## Methods

### Ethics and clinical stratification

Total blood of healthy donors in the United Kingdom was obtained following approval by the Southwest - Cornwall & Plymouth Research Ethics Committee (REC 23/SW/0029).

Total blood from PLHTLV and healthy donors in Brazil was obtained following approval by the Research Ethics Committees of each reference center and by CONEP (National Commission for Research Ethics), under approval number 7.742.287. All patients gave written informed consent before blood collection in accordance with the Declaration of Helsinki protocol. The diagnosis of HTLV-1 infection was based on the availability of previous serological screening (chemiluminescence ELISA) and confirmatory PCR or Western blot. The stratification of ATL cases followed the Shimoyama classification^52^. The diagnosis of HAM/TSP was based on clinical evaluation and laboratory testing, with spinal MRI and neurophysiological studies used to support the diagnosis and help differentiate from other neurological conditions.

### Cell lines

HTLV-1 infected (MT-2, MT-4, OS-P2, C10/MJ, C8166-45) and uninfected (Jurkat E6.1, CEM-ss, HuT-78, MOLT-4, Sup-T1, U937) cell lines were obtained through the Biodefense and Emerging Infections Research (BEI) Resources Repository (Catalogue Numbers: ARP-237, ARP-120, ARP-1393, ARP-4407, ARP-404, ARP-776, ARP-89, ARP-175, ARP-100) except for U937 cells, which were a kind gift from Dr. Andrea Savarino (Italian Institute of Health) and Jurkat E6.1 cells, which were obtained from the European Collection of Authenticated Cell Cultures (ECACC Catalogue No. 88042803). ATL cell lines (MT-1, SP, TL-Om1, ATL-55T(+), ED) were a kind gift of Dr. Jun-ichirou Yasunaga (Kumamoto University).

Cells were cultured in RPMI medium (Merck) complemented with 10% FCS and 1% penicillin/streptomycin. The medium of SP and ATL-55T(+) cells was also complemented with 100 U/mL IL-2.

HEK293-T cells (ECACC Catalogue No. 12022001) were cultured in DMEM medium complemented with 10% FCS and 1% penicillin/streptomycin.

Cells were maintained at 37 °C in a 5% CO_2_ incubator and periodically tested to exclude mycoplasma contamination.

### Primary cells

Peripheral blood mononuclear cells (PBMCs) were isolated within 4h of blood collection using Ficoll-Paque PLUS (Cytiva). Briefly, the blood was diluted 1:1 in PBS and layered over Ficoll at a ratio of 2:1. After centrifugation at 400 x g for 30 min, the PBMC layer was transferred to a clean tube and washed twice with PBS. Cell viability was assessed using trypan blue exclusion and quantified with an automated Countess II cell counter (Invitrogen). Immediately after isolation, PBMCs were cryopreserved and stored in liquid nitrogen until further use.

CD4 T cells were isolated either directly from total blood using the RosetteSep™ Human CD4+ T Cell Enrichment Cocktail (Stemcell) or from frozen PBMCs using the EasySep™ Human CD4+ T Cell Isolation Kit (Stemcell) according to manufacturer’s instruction. Cells were cultured in RPMI medium (Merck) complemented with 10% FCS, 1% penicillin/streptomycin and 20-100 U/mL IL-2. To compare resting and activated uninfected cells, CD4 T cells were stimulated by adding phytohemagglutinin (PHA; Gibco) at a 1:100 dilution to the complete culture medium.

### TMT Proteomics

HTLV-1^+^/ATL (MT-2, MT-4, SP) and uninfected (Jurkat, CEM-ss, MOLT-4) cell lines were lysed in RIPA buffer complemented with Protease and Phosphatase Inhibitor Cocktail (cOmplete Protease Inhibitor Cocktail and PhosSTOP, Roche). Lysates were prepared in quadruplicate for each cell line. Lysate concentration was quantified by Pierce BCA assay (Thermo Fisher Scientific) and confirmed as >2mg/mL for all samples.

Protein samples (100 µg each) were digested with trypsin (2.5 µg per 100 µg protein; 37 °C, overnight), labelled with TMTpro reagents according to the manufacturer’s protocol (Thermo Fisher Scientific), and pooled. An aliquot of 100 µg of the pooled sample was desalted using a SepPak cartridge according to the manufacturer’s instructions (Waters, Milford, Massachusetts, USA). Eluate from the SepPak cartridge was evaporated to dryness and resuspended in buffer A (20 mM ammonium hydroxide, pH 10) prior to fractionation by high pH reversed-phase chromatography using an Ultimate 3000 liquid chromatography system (Thermo Fisher Scientific). In brief, the sample was loaded onto an XBridge BEH C18 Column (130Å, 3.5 µm, 2.1 mm X 150 mm, Waters, UK) in buffer A and peptides eluted with an increasing gradient of buffer B (20 mM Ammonium Hydroxide in acetonitrile, pH 10) from 0-95% over 60 minutes. The resulting fractions (concatenated into 20 in total) were evaporated to dryness and resuspended in 1% formic acid prior to analysis by nano-LC MS/MS using an Orbitrap Fusion Lumos mass spectrometer (Thermo Scientific). High pH RP fractions were further analyzed using an Ultimate 3000 nano-LC system in line with an Orbitrap Fusion Lumos mass spectrometer (Thermo Scientific). Briefly, peptides in 1% (vol/vol) formic acid were injected onto an Acclaim PepMap C18 nano-trap column (Thermo Scientific). After washing with 0.5% (vol/vol) acetonitrile 0.1% (vol/vol) formic acid peptides were resolved on a 500 mm × 75 μm Acclaim PepMap C18 reverse phase analytical column (Thermo Scientific) over a 150 min organic gradient, using 7 gradient segments (1-6% solventB over 1min., 6-15%B over 58min., 15-32%B over 58 min., 32-40%B over 5 min., 40-90%B over 1min., held at 90%B for 6min and then reduced to 1%B over 1 min.) with a flow rate of 300 nl min−1. Solvent A was 0.1% formic acid and Solvent B was aqueous 80% acetonitrile in 0.1% formic acid. Peptides were ionized by nano-electrospray ionization at 2.0kV using a stainless-steel emitter with an internal diameter of 30 μm (Thermo Scientific) and a capillary temperature of 300°C. All spectra were acquired using an Orbitrap Fusion Lumos mass spectrometer controlled by Xcalibur 3.0 software (Thermo Scientific) and operated in data-dependent acquisition mode using an SPS-MS3 workflow. FTMS1 spectra were collected at a resolution of 120000, with an automatic gain control (AGC) target of 200000 and a max injection time of 50ms. Precursors were filtered with an intensity threshold of 5000, according to charge state (to include charge states 2-7) and with monoisotopic peak determination set to Peptide. Previously interrogated precursors were excluded using a dynamic window (60s +/-10ppm). The MS2 precursors were isolated with a quadrupole isolation window of 0.7m/z. ITMS2 spectra were collected with an AGC target of 10 000, max injection time of 70ms and CID collision energy of 35%. For FTMS3 analysis, the Orbitrap was operated at 50000 resolution with an AGC target of 50000 and a max injection time of 105ms. Precursors were fragmented by high energy collision dissociation (HCD) at a normalised collision energy of 60% to ensure maximal TMT reporter ion yield. Synchronous Precursor Selection (SPS) was enabled to include up to 10 MS2 fragment ions in the FTMS3 scan.

Raw data files were processed and quantified using Proteome Discoverer software v2.4 (Thermo Scientific) and searched against the UniProt Human database (downloaded January 2024: 82415 entries), the Uniprot HTLV-1 database (downloaded June 2024: 9 entries) and the P13 sequence using the SEQUEST HT algorithm. Peptide precursor mass tolerance was set at 10ppm, and MS/MS tolerance was set at 0.6Da. Search criteria included oxidation of methionine (+15.995Da), acetylation of the protein N-terminus (+42.011Da) and methionine loss plus acetylation of the protein N-terminus (−89.03Da) as variable modifications and carbamidomethylation of cysteine (+57.0214) and the addition of the TMTpro mass tag (+304.207) to peptide N-termini and lysine as fixed modifications. Searches were performed with full tryptic digestion and a maximum of 2 missed cleavages were allowed. The reverse database search option was enabled, and all data was filtered to satisfy false discovery rate (FDR) of 5%.

Protein grouping and total peptide-amount normalisation were performed in Proteome Discoverer 2.4, while master-protein refinement and all subsequent statistical analyses were conducted in R (v4.4.0). The master-protein improvement script used queried UniProt to obtain the current annotation status for all protein accessions (updated 2024-07-17), updating redirected or obsolete entries accordingly, and then selecting the best-annotated UniProt candidate, prioritising reviewed records. Abundance values were Log2 transformed and used as input for pathway analysis with the Camera gene set algorithm in ReactomeGSA (Reactome Version 93)^53^. Differential expression analysis was performed using the limma package (v3.64.3)^54^ in Rstudio. Differential expression results were filtered using the Mitocarta database (v 3.0) ^22^.

### Metabolomic analysis

The analysis of intracellular metabolic extracts from HTLV-1^+^/ATL (MT-2, MT-4, SP) and uninfected (Jurkat E6.1, MOLT-4, CEM-ss) cell lines was conducted as previously described^21^. Briefly, a cold extraction solution containing 80% methanol and 20% water was used to extract polar metabolites. Samples were analyzed by LC-MS/MS on a Thermo Scientific Q Exactive Plus mass spectrometer coupled to a Vanquish UHPLC System. Liquid chromatography separation was performed under HILIC pH 9 condition using a ZIC-pHILIC column. Peak areas, representing metabolite levels, were extracted using Thermo Compound Discoverer 3.3 SP3. Only metabolites identified either by matching their retention time and fragmentation spectra to that of authentic pure standards or by spectral matching to compounds in the mzCloud database were retained for downstream analysis. One quality control (QC) sample pool was made by pooling a small aliquot from each sample. This pool was analyzed at the beginning, middle, and end of the sample sequence. Samples were analyzed in a pseudorandomized order, with 4 µL injected per run. Metabolite abundances were normalized against both QC samples and the total metabolite signal to account for differences in sample amount. The resulting normalized values were Log_10_ transformed and used as input for downstream analysis in MetaboAnalyst^55^ (v6.0). Enrichment analysis was conducted using the dedicated module in MetaboAnalyst with The Small Molecule Pathway Database (SMPDB) metabolite set serving as reference library and including only metabolite sets containing at least two metabolites. Hierarchical clustering analysis was performed in the One-factor Statistical Analysis module of MetaboAnalyst. The data were row-scaled, and clustering was performed using Euclidean distance as the similarity metric and the Ward linkage method for cluster aggregation. The top twenty metabolites, ranked by t-test results after Benjamini–Hochberg multiple-testing correction, were selected for visualization in the heatmap.

### Mitochondrial isolation

Mitochondria were isolated using the Mitochondria Isolation Kit, human (Miltenyi Biotec) following manufacturer’s instructions. Briefly, 20 x 10^6^ cells were washed twice with PBS and then lysed with lysis buffer complemented with cOmplete Protease Inhibitor Cocktail (Roche). Cells were homogenized using a cell homogenizer (isobiotec) with a 10µm clearance ball and 8-10 strokes. The homogenized lysate was then incubated with anti-TOM22 MicroBeads for 30 min at 4 °C. Mitochondria were purified by magnetic isolation using LS Columns (Miltenyi Biotec).

### Western Blot

Whole cells or isolated mitochondria were lysed in RIPA buffer complemented with cOmplete Protease Inhibitor Cocktail (Roche) for 5 minutes at 4 °C. Protein lysates were cleared by centrifugation at 8,000 × g for 10 min and either used immediately or snap-frozen in liquid nitrogen and stored at −80 °C for subsequent use. For electrophoresis, protein lysates were mixed with mPAGE loading buffer (Merck) and β-mercaptoethanol, then incubated at 70 °C for 10 min. Samples were run on a 4–12% gradient gel at 100–150 V and proteins were transferred onto a nitrocellulose membrane at 100 V for 1h at RT. Membranes were blocked with PBS Blocking Buffer (LICORbio) for 1h at RT, followed by overnight primary antibody incubation at 4 °C. The primary antibodies used included anti-FDXR (1:1000, Rabbit, Proteintech #15584-1-AP), anti-p53 (Rabbit, Cell Signaling #2527), anti-p63α (1:1000, Rabbit, Cell Signaling #13109), anti-p73 (1:1000, Rabbit, Cell Signalling #14620), anti-S. pyogenes-Cas9 (1:1000, Mouse, Cell Signalling #14697) anti-VDAC (1:500, Rabbit, Invitrogen #MA5-33205), anti-Tax (1:250, Mouse, Merck #MABF3063), anti-HTLV-1-p24 (1:250, Mouse, Santa-Cruz #sc-53891), anti-HTLV-1-gp46 (1:250, Mouse #sc-53890), anti-alpha Tubulin (1:15000, Rabbit, GeneTex #GTX112141), anti-beta Actin (1:15000, Mouse, Cell Signaling #3700). After three washes in PBS 0.1% tween, membranes were incubated with secondary antibodies coupled with IRDye® 680 or 800 for 1h RT in the dark. Membranes were imaged using an Odyssey XF imaging system (LICORbio). Quantifications were performed with the Empiria Studio software (v3.2, LICORbio).

### Immunofluorescence

Cells were seeded on poly-D-lysine–coated 18-well chamber slides (Ibidi). To visualize mitochondria, cells were stained with Mitotracker Red CMXRos Dye (1:500, Invitrogen #M46752) for 30min at 37 °C. The Click-iT™ EdU Cell Proliferation Kit (ThermoFisher) was used to stain proliferating primary CD4 T cells, according to manufacturer’s instructions. To this purpose cells were first incubated with 10 µM Edu for 2h at 37 °C. Cells were then fixed with 4% paraformaldehyde (PFA) for 1 h at RT, followed by permeabilization with 0.1% Triton X-100 in PBS for 10 min at RT. EdU incorporation was detected using the Click-iT reaction cocktail as provided in the kit. For subsequent immunofluorescent staining, cells were blocked with 1% BSA in PBS for 30 min at RT and primary antibody incubation (anti-FDXR, 1:100, Rabbit, Proteintech #15584-1-AP) was performed in a humidified chamber at 37 °C for 1h. Secondary antibody (Goat anti-Rabbit IgG, Alexa Fluor^TM^ 488 or 647, 1:3000 in blocking buffer) was incubated together with DAPI (1:3000) in a humidified chamber at 37 °C for 30 minutes. Wells were filled with Ibidi mounting medium, and samples were imaged using a Leica SP8 confocal microscope. Images were analysed with Fiji (v 2.16) and Imaris (v11, Oxford Instruments).

### Transcriptomic data reanalysis

RNA-Seq data of cell lines were retrieved from Protein Atlas. Normalized transcripts per million (nTPM) data for FDXR expression were extracted and filtered to retain only data from blood cancer (n= 3 HTLV^+^, 93 leukemia, 73 lymphoma, 34 myeloma) and non-cancerous (n= 63) cell lines.

Patient-derived microarray data were retrieved from the Gene Expression Omnibus (GEO) repository. Studies were identified using the keyword “HTLV-1” and included based on the following criteria: (1) availability of primary CD4 T cells from both PLHTLV and healthy donors; (2) use of whole-genome microarray platforms; (3) stratification of samples by disease status; and (4) inclusion of at least two donors per condition. Filtering with these criteria yielded four datasets that were used for downstream analysis: GSE33615^12^, GSE38537^25^, GSE132666^26^, GSE233437^27^.

Raw data were background corrected and normalised using the normexp and quantile functions of limma^54^. Probes were annotated to gene symbols using platform-specific annotation databases, and expression values for multiple probes mapping to the same gene were averaged. Samples corresponding to PBMCs or CD4-negative populations, as well as samples with unknown ATL subtype status, were excluded prior to integration. Batch effects between studies were corrected using the ComBat function of the sva package^56^ with each GEO study being defined as a separate batch.

Bulk RNA-Seq data obtained from samples of ATL (n=57), asymptomatic HTLV-1 (n=3) and healthy controls (n=3) were retrieved from the European Genome-phenome Archive (accession EGAD00001001411, ref^8^). One ATL sample lacked subtype annotation and was excluded from downstream analyses. Samtools^57^ (v1.22) was used to sort bam files and convert them to paired end-fastq files. Adapter sequences were removed and low-quality bases trimmed with fastp^58^ (v0.23.4), and the resulting reads were aligned to the human reference genome (GRCh38) using STAR (v2.7.11b)^59^. Gene-level read counts were obtained with the featureCounts function^60^ (v2.0.8) of the Subread suite. Differential gene-expression analysis was performed with DESeq2^61^ (v1.48.2). Normalized *FDXR* expression values for plotting were derived by applying the variance-stabilizing transformation implemented in DESeq2.

For single cell RNA-seq, raw sequencing data were downloaded from Short Read Archive under study accession PRJEB47382^39^. A custom reference transcriptome was obtained merging human GRCh38 annotation (10x Cell Ranger pre-built reference version 2024-A) with Human T-cell leukemia virus type I genome available at GenBank (accession and version AB513134.1). For all samples, raw reads were aligned to the custom reference transcriptome and UMI counts calculated using the 10x Cell Ranger Count pipeline (version 9.0.0). The expression data were imported in R (version 4.1.1) and analyzed with the Seurat^62^ (version 4.3.0) and popsicleR^63^ (version 0.2.1) R packages. For each sample, low quality cells were identified as outliers based on the number of genes, UMI counts, and the percentage of reads mapping to mitochondrial genes per cell, and these were subsequently discarded. Doublets were detected and removed using the scDblFinder^64^ (version 1.8.0) R package. Cell cycle scores were assigned to each cell using Seurat, with automated identification of cells in S or in G2/M phases of the cell cycle. Principal component analysis (PCA) was used to reduce dataset dimensionality; Louvain clustering and UMAP visualization were performed on PCA reduced embedding after selecting an appropriate number of principal components for each sample. Upon the evaluation of an optimal clustering resolution, BlueprintEncode reference dataset from SingleR^65^ (version 1.8.1) and known marker genes were used to annotate clusters as T/NK cell, B cell, myeloid and contaminants (e.g., platelets, erythrocytes). Cells annotated as T/NK lymphocytes (94,834 cells in 17 samples) were integrated using the Seurat integration strategy (FindIntegrationAnchors and IntegrateData functions) and re-clustered with the same procedure described above. The T and NK cell clusters were classified monitoring the expression of known markers. Low-quality and contaminated clusters were filtered out after inspecting the number of genes, UMI counts and doublet score, resulting in 10 clusters comprising 91,436 good quality cells, of which 5 clusters (65,319 cells) are annotated as CD4 T cells.

### Survival and prognostic analysis

To evaluate the prognostic relevance of *FDXR* expression, we used the RNA-seq data from 51 patients of dataset EGAD00001001411(ref ^8^) for which a clinical follow-up was available^8,66^. Overall survival was censored at hematopoietic stem cell transplantation. Cox proportional-hazards models using continuous Z-score values of normalized *FDXR* expression were applied to the entire cohort as well as in separate analysis for the aggressive (acute and lymphoma) and indolent (chronic and smoldering) ATL subtypes. The proportional-hazards assumption was verified using Schoenfeld residuals both in the overall cohort and in each subtype. For Kaplan–Meier analyses, *FDXR* Z-scores were used to stratify patients into either two or four groups based on SD thresholds, and survival differences were evaluated using the log-rank test.

### In Silico Inference of Transcription Factor Activity

The RNA-Seq data of all leukemia/lymphoma cell lines retrieved from Protein Atlas were stratified based on nTPM *FDXR* expression (mid-high FDXR group: ≥ 75 nTPM; mid-low FDXR group: < 75 nTPM).

The Discriminant Regulon Expression Analysis (DoRothEA) package^30^ was used to obtain high- to medium-confidence (A–C) human transcription factor (TF)–target interactions. Gene expression values (nTPM) were Log2 normalized and a context-specific regulatory network was built using the 1,000 most variable genes across all RNA-Seq samples, together with all TFs. Expression values for each gene were discretised into three equal-frequency bins using the infotheo (v 1.2.0.1) package. The minet (v 3.66.0) package^31^ was used to infer regulatory interactions by first computing a mutual information matrix from discretised expression data (build.mim, estimator = “mi.empirical”) and then applying the Algorithm for the Reconstruction of Accurate Cellular Networks (ARACNe) for network pruning (ε = 0.05). To obtain the final regulon, the resulting unsigned network was overlapped with DoRothEA to assign regulatory direction (activation vs repression) where available. The signed network was formatted into a regulon object where each TF is linked to its targets with a mode of regulation (tfmode) and an associated likelihood score. The Virtual Inference of Protein-activity by Enriched Regulon analysis (VIPER) algorithm^29^, was then applied (scaling method: median absolute deviation) to the Log2 normalized expression matrix to infer sample-specific TF activity scores. The statistical significance of differences in TF activity between the mid–high and mid–low FDXR groups was evaluated using the limma (v3.64.3) package^54^, as outlined in the Statistical Analysis section.

Upstream TF regulators of TP63 and TP73 were extracted from the final, signed, regulon. For each upstream TF, the regulatory weight (mor) was plotted against its differential RNA expression between the mid-high and mid-low FDXR groups, computed using the limma package. Plots were generated using the ggplot2 (v4.0.0) package in Rstudio.

### Chromatin immunoprecipitation (ChIP)-qPCR

ChIP was performed using the SimpleChIP® Enzymatic Chromatin IP Kit (Cell Signaling Technology), according to manufacturer’s instructions. Briefly, 10 × 10^6^ HTLV-1–infected (C8166-45, MT-4), ATL (SP), or uninfected (Jurkat, MOLT-4) cells were resuspended at 0.5 × 10^6^ cells/mL and cross-linked with 1% formaldehyde. After 20 min, cross-linking was quenched by the addition of 10X glycine, and cells were washed twice with ice-cold PBS. To isolate nuclei, cells were lysed sequentially in Buffer A and Buffer B supplemented with protease inhibitor cocktail and dithiothreitol (DTT). Isolated nuclei were digested using micrococcal nuclease for 20 min at 37 °C. Nuclei were pelleted and resuspended in 1X ChIP buffer containing protease inhibitor cocktail, followed by sonication (60% amplitude, 3 cycles of 20 s, with 30 s on ice between cycles) using a QSonica sonicator. Lysates were clarified by centrifugation, and an aliquot was used to confirm chromatin fragment size and quality by agarose gel electrophoresis. For immunoprecipitation, 5–10 µg of chromatin was incubated with anti-p63α Rabbit mAb (Cell Signaling #13109), anti-ΔNp63 (Cell Signaling #67825S), anti-Histone H3 Rabbit mAb (Cell Signaling #4620), or rabbit IgG control (Cell Signaling #8726) overnight at 4 °C with rotation. Immune complexes were incubated with protein G magnetic beads for 2 h at 4 °C, followed by three low-salt washes and one high-salt wash (5 min each at 4 °C). Chromatin was eluted using SDS-containing ChIP elution buffer for 30 min at 65 °C, and cross-links were reversed by incubation with NaCl and proteinase K for 2 h at 65°C. DNA was purified using spin columns provided in the kit. qPCR was performed using primers specific for the p53 binding element in the FDXR promoter, as previously described.^33^. PCR conditions were as follows: an initial denaturation at 95°C for 5 min, followed by 40 cycles of 95 °C for 30 s, 62 °C for 30 s, and 72 °C for 30 s, with a final extension at 72 °C for 5 min. Primer sequences are listed in Supplementary Table 4. Following amplification, 10 µL of PCR product was analyzed by 2% agarose gel electrophoresis.

### CHIP-Seq data analysis

Previously published ChIP-seq datasets for IRF4, BATF3, H3K27ac, and BRD4 from ATL^35^ and control cells were retrieved from ChIP-Atlas (v3.0)^67^ using the following SRX identifiers: SRX2548277, SRX2548278, SRX2548279, SRX2548280, SRX2548281, SRX2548282, SRX2548284, SRX2548286, SRX2548291, SRX2548292, SRX2548295, SRX2548296, SRX2548299, SRX2548300, SRX1528530, SRX398094. For each dataset, the standardized ChIP-Atlas processed files were used, which include RPM-normalized BigWig coverage tracks and MACS2 peak calls (BED4 format; hg38; q = 1e-05). ChIP-Atlas 3.0 processing follows a uniform pipeline using Bowtie2 (v2.2.2) for alignment, Bedtools (v2.17.0) for normalized coverage calculation, and MACS2 (v2.1.0) for peak calling. All tracks were visualized using the Integrative Genomics Viewer^68^ (IGV), together with annotation files obtained from ChIP-Atlas and IGV.

AICE^35,48^ motif occurrences within the TP63 genomic region and overlapping with IRF4 and BATF3 peaks in ATL cells, were detected performing motif scanning with the Hypergeometric Optimization of Motif EnRichment (HOMER^69^, v3.12). ChIP-seq peaks (threshold q = 1e-05) located within the genomic interval chr3:189,627,904–189,898,305 (hg38) were extracted in IGV and exported as a BED file. Motif analysis was conducted with the annotatePeaks.pl and findMotifsGenome.pl utilities in HOMER, using a custom motif file containing three previously described^48^ canonical AICE motifs (GAAATGAnTCA, TGAnTCAGAAA, TTTCnnnnTGAnTCA). For each BATF3 or IRF4 peak, a 100 bp window centered on the peak summit was extracted and used for motif scanning after masking repetitive sequences. The motif-peak distances were plotted using the ggplot2 (v4.0.0) package in Rstudio.

### RNA and DNA isolation

RNA and genomic DNA were extracted from cell lines and primary cells using the AllPrep DNA/RNA Isolation Kit (Qiagen) according to the manufacturer’s instructions. Prior to nucleic acid isolation, cell lysates were homogenized using QIAshredder columns and centrifuged for 2 min at 17,000 × g. RNA was eluted in 30 µL of nuclease-free water, and genomic DNA was eluted in 50 µL of EB buffer.

### Gene expression quantification by qPCR

Isolated RNA was reverse transcribed using the iScript Reverse Transcription Supermix (Bio Rad) according to manufacturer’s instructions. Relative viral and host gene expression was analysed by duplex qPCR using the primers and probes listed in Supplementary Table 4. The TATA-box binding protein (TBP) gene was used as the housekeeping control to normalize expression levels. PCR reactions were carried out in a CFX Opus 96 machine (Bio-Rad) under the following cycling conditions: an initial denaturation at 95 °C for 30 sec, followed by 49 cycles of denaturation at 95 °C for 5 seconds, annealing-extension at 56 °C for 30 seconds. Data were analysed using Biorad CFX Maestro Software v2.3. For each gene of interest (GOI) and sample, the relative quantity (RQ) was calculated as: E_GOI_ ^ (Cq(min) − Cq(sample)), where E = is the efficiency of the primer and probe set calculated with the formula: (% Efficiency × 0.01) + 1, Cq(min) is the average Cq for the sample with the lowest average Cq, and Cq(sample) is the average Cq value for the sample of interest. The RQ values were then used to obtain the normalized expression (ΔΔCq) levels for each GOI, calculated as: RQ(sample, GOI) ÷ RQ(sample, TBP).

### Proviral load quantification by digital droplet PCR (ddPCR)

HTLV-1 proviral load was measured by ddPCR as previously described^70^. Primer and probe sequences are listed in Supplementary Table 4. Genomic DNA was extracted from primary CD4 T cells, and proviral load was calculated as: PVL(%) = (HTLV-1 pX copy nr) / (albumin copy nr / 2) x 100.

### Gene expression knockout with CRISPR/Cas9

HTLV-1 infected (MT-4) and ATL (ATL-55T) cell lines were transduced to stably express Cas9 using the lentiCas9-Blast lentivirus (Addgene #52962-LV)^71^ at MOI 50. Transduced cells were selected and kept in selection with blasticidin (5 µg/mL) and the expression of Cas9 was confirmed by western blot in each experiment. The sgRNA sequences (listed in Supplementary Table 4) targeting TP63 and TP73 were retrieved from the Brunello library^72^ and oligos were synthesized (Eurofins Genomics) with the overhangs required for cloning in the lentiGuide-Puro vector (Addgene #52963)^71^. Cloning was performed following the standardised protocol of the Zhang laboratory^71^. A plasmid encoding for non-targeting sgRNA cloned in lentiGuide-Puro was used as control (Addgene #80262). Plasmids containing cloned sgRNAs were co-transfected with the packaging plasmid psPAX2 (AddGene #12260) and the envelope plasmid pVSVg (AddGene #8454) at a ratio of 4:3:1 in HEK293-T cells. The lentiviruses obtained were used to transduce stably Cas9 expressing cells. Transduced cells were selected with puromycin (2 µg/mL) for two weeks and both Cas9 expression and knockout efficiency were confirmed by western blot.

### In silico drug efficacy prediction

The *in silico* screening of candidate compounds for cytotoxicity assessment was performed on data generated by the Cancer Target Discovery and Development (CTD²) Network (https://www.cancer.gov/ccg/research/functional-genomics/ctd2) established by the National Cancer Institute’s Center for Cancer Genomics. Specifically, the AUC values for individual compounds were obtained from the dataset originally described in^73^. The correlation between *FDXR* mRNA expression levels and drug efficacy was analyzed using the methodology outlined in^41^ with significance thresholds determined using a Bonferroni-adjusted two-tailed z-test.

### Drug Treatments

*In vitro* and *ex vivo* treatments were performed for 48–72 h, as specified in the corresponding figure legends, using fluorouracil (Cambridge Bioscience) or bleomycin sulfate (Merck). All compounds were dissolved in DMSO. Corresponding serial dilutions of DMSO were used as matched controls in drug sensitivity assays.

### MTS assay

Cell viability/proliferation was assessed with the MTS assay (Promega), according to manufacturer’s instruction. Briefly, 0.5 × 10^4^ cells were plated in triplicate into a flat-bottom 96-well plate, and 20 µL of MTS + phenazine methosulfate (PMS) solution was added to each well at the end of the drug incubation period. Cells were kept 1–3 h at 37 °C and 5% CO_2_ and the 490nm absorbance was measured using a SPECTROstar Nano (Bmg Labtech) microplate reader.

### Drug-Seq

CD4 T cells of five ATL patients, three healthy donors and two ATL cell lines (SP and TLOm-1) were treated for 48h with fluorouracil, bleomycin sulfate (5 µM or 10 µM of each drug), or DMSO control. Total RNA was isolated and used for library preparation and sequencing using a highly multiplexed 3′-end bulk RNA barcoding and sequencing approach (MERCURIUS™ BRB-seq service, Alithea Genomics SA)^74^. RNA quantity and quality were assessed using a NanoDrop spectrophotometer (Thermo Fisher Scientific, USA), and RNA concentrations were normalized to 100 ng/µL. Bulk RNA barcoding and sequencing (BRB-seq) libraries were generated using the MERCURIUS™ BRB-seq Library Preparation Kit for Illumina (Alithea Genomics, #10813) according to the manufacturer’s instructions. Libraries were sequenced on an AVITI platform (Element Biosciences). Raw sequencing reads were processed using STAR (v2.7.9a) in STARsolo mode, enabling simultaneous alignment, barcode demultiplexing, and gene quantification. Barcode demultiplexing allowed one mismatch (--soloCBmatchWLtype 1MM), and UMI deduplication was performed using the 1MM_Directional method (--soloUMIdedup 1MM_Directional). Reads were aligned to the human reference genome GRCh38, and gene-level quantification was performed using --quantMode GeneCounts, generating raw and UMI-deduplicated count matrices. UMI-deduplicated counts were used for downstream analyses to correct for PCR amplification bias. Differential expression was performed in DESeq2 (v1.50.2) using interaction designs (Condition×DrugDose or Group×DrugDose, where Condition encodes disease status and Group encodes disease status and cell type). Wald statistics were used to rank genes. Preranked GSEA was run with clusterProfiler (v4.18.4) against MSigDB Hallmark gene sets (obtained via the msigdbr package, v25.1.1), using the conventional^75^ GSEA FDR < 0.25 threshold. Enrichment curves and ridge plots were generated with the enrichplot (v1.30.4) and ggridges (v0.5.7) packages, respectively.

### Statistical analysis

The significance of HTLV-1^+^ cell line ranking by *FDXR* expression among the Human Protein Atlas blood cell lines was assessed using a permutation-based test (100,000 permutations). In each permutation, FDXR expression values were randomly reassigned across cell lines and the number of HTLV-1^+^ cell lines ranked among the top five expressers was recorded. The empirical p-value was defined as the proportion of permutations yielding an enrichment equal to or greater than that observed in the original dataset.

The significance of differences in log₂ *FDXR* expression across disease conditions in the meta-analysis of microarray datasets was assessed using a linear mixed-effects model fitted with the lme4 (v1.1.38) package, with disease status included as a fixed effect and the study from which each dataset was derived included as a random intercept. Post hoc contrasts comparing each disease group with uninfected controls were estimated from study-adjusted marginal means using the emmeans (v2.0.1) package, with p-values adjusted for multiple testing using the Holm method. Differences in relative qPCR gene expression were assessed using log2-transformed values and analyzed by unpaired Wilcoxon rank-sum test for two-group comparisons, or by Kruskal–Wallis test followed by Dunn’s post hoc test for multiple-group comparisons. Differential TF activity between mid-high and mid-low FDXR groups was assessed using the limma package, applying empirical Bayes–moderated linear models with contrasts to obtain FDR-adjusted p-values.

Drug inhibition curves were fitted for each cell line using a four-parameter nonlinear regression (4PL) and the fitted curves were used to compute AUCs using the MESS package (v0.6.0). Differences in AUC values between HTLV-1^+^/ATL and uninfected cell lines were assessed using an unpaired Wilcoxon rank-sum test.

The effects of Fluorouracil and Bleomycin on cell proliferation (EdU incorporation status) were analysed at the single-cell level using a binomial mixed-effects model fitted with the lme4 package, with treatment as a fixed effect and donor as a random intercept. Estimated marginal means were computed using the emmeans package, and Dunnett-adjusted post hoc comparisons were performed for each treatment versus the DMSO control.

## Data availability

Proteomics data generated in this study have been deposited in the ProteomeXchange Consortium via PRIDE (accession number PXD073455). All other data supporting this study are available in the article and its Supplementary Information or from the corresponding author upon reasonable request.

## Author contributions

Conceptualization, ILS; Methodology, LG, SOGM, ILS; Investigation (wet-lab experiments), LG, MJ, AARS, VCF, MESA, TR, PK, GLDAB, RASS, LAL, AL, BH, FCCS, SS, AQ, SLMG; Investigation (proteomics) PAL, KH; Investigation (metabolomics) WZ, ARG; Formal Analysis (experimental data), LG, ILS; Formal Analysis (bioinformatic data), YK, PAL, MF, ILS; Resources, LG, FEL, SO, KK, SOGM, ILS; Cohort Recruitment and Clinical Management, YN, JP, HFC, TK, KS, ECS, FEL, SOGM; Writing – Original Draft, ILS, LG; Writing – Review & Editing, all authors; Supervision, LG, ECS, FEL, KK, SOGM, ILS; Funding Acquisition, LG, SOGM, ILS.

## Acknowledgments

The authors are thankful to the individuals who donated blood for the purpose of this study. The authors thank Dr. Jun-ichirou Yasunaga (Kumamoto University) for sharing MT-1, ED, TL-Om1 and ATL-55T cells, Dr. Andrea Savarino (Italian Institute of Health) for sharing U937 cells, and the Biodefense and Emerging Infections Research (BEI) Resources Repository for providing MT-2, MT-4, C10/MJ, OS-P2, SP, CEM-ss, Sup-T1 and MOLT-4 cells. The authors would like to acknowledge the Wolfson bioimaging facility (University of Bristol) for microscopy support. ILS and SOG acknowledge funding from Royal Society (RGS\R1\231061, ILS; and ICAO\R1\231054 ILS and SOG). ILS and LG, acknowledge funding from the Marie Skłodowska-Curie Postdoctoral Fellowship – UKRI Funding guarantee (EP/Z002850/1).

## Supplementary Figures

**Supplementary Figure 1. Mitochondrial metabolic alterations in HTLV-1^+^/ATL cell lines.**

**(A)** Experimental design. The metabolome and proteome of three HTLV-1^+^/ATL cell lines (MT-2, MT-4, SP) were compared with three uninfected lymphoid cell lines (Jurkat, MOLT-4, CEM-ss). Cell lysates were analysed in triplicate for metabolomics and in quadruplicate for proteomics.

**(B)** Pathway enrichment analysis of metabolomic profiles comparing HTLV-1^+^ and uninfected cells.

**(C)** Heatmap of the twenty most significantly altered metabolites in HTLV-1^+^ versus uninfected cells, ranked by t-test followed by Benjamini–Hochberg multiple-testing correction.

**(D)** Proteomic pathway analysis showing the ten most upregulated pathways in HTLV-1–infected cells.

Metabolomic data were analysed with MetaboAnalyst using SMPDB metabolite sets. Proteomic pathway analysis was performed using CAMERA gene set analysis with Reactome gene sets. Pathways that are exclusively or predominantly mitochondrial in panels (B) and (D) are highlighted in red, and metabolites from these pathways that appear in the heatmap (C) are indicated by asterisks.

**Supplementary Figure 2. Relative HTLV-1 RNA and protein levels in the cell models used in the study.**

**(A-C)** Relative expression of the HTLV-1 transcripts *Hbz* (A), *Gag/Pol* (B) and *Tax/Rex* (C) were measured by qPCR using the primers listed in Supplementary Table 4. Each dot represents an independent qPCR assay, and data are presented as mean ± SD. Relative RNA levels were calculated by ΔΔCq using *TBP* expression as reference.

**(D-E)** Protein levels of HTLV-1 Env gp46 and p24 (D) and Tax (E) were assessed by western blot. The fusion protein Tax-Env, typical of the MT-2 cell line, is visible in both (D), as detected with an anti-gp46 antibody, and in (E) as detected with an anti-Tax antibody.

**Supplementary Figure 3. *FDXR* mRNA levels in PLHTLV and ATL survival curves by quartiles of FDXR expression.**

(**A**) CD4 T cells were isolated from PBMCs obtained from the cohort described in Supplementary Table 3.

(**B-C**) Relative *FDXR* mRNA levels were calculated by ΔΔCq using *TBP* expression as reference. Each dot represents an individual donor, while black lines depict medians of each subgroup. In panel (B) differences in relative *FDXR* expression between PLHTLV and healthy controls were analyzed by unpaired Wilcoxon rank-sum test following log_2_ transformation.

(**D**) Kaplan–Meier analysis of overall survival in patients grouped into quartiles based on the median Z-score of *FDXR* expression. *FDXR* expression values were derived from RNA-seq dataset EGAD00001001411 and analysed in ATL patients with available survival data (n = 51^8,66^). Overall survival was censored at hematopoietic stem cell transplantation, and survival differences were assessed using the log-rank test. *p <0.05

**Supplementary Figure 4. p53 family expression profiles and p63 chromatin binding at the FDXR promoter in HTLV-1^+^/ATL and uninfected cells.**

**(A)** Isoform-specific primer design for amplification of TA and ΔN isoforms of TP63 and TP73 by qPCR.

**(B,C)** Relative mRNA expression of ΔN isoforms of TP63 (B) and TP73 (C) in HTLV-1^+^, ATL, and uninfected cell lines and primary cells, as assessed by qPCR. In (B), primary keratinocytes were used as a positive control for ΔNTP63 expression.

**(D,E)** Relative mRNA (D) and protein (E) expression of p53 in HTLV-1^+^, ATL, and uninfected cell lines and primary cells, as assessed by qPCR and western blotting, respectively.

In panels B-D, each dot represents an independent qPCR assay (n = 3), and data are presented as mean ± SD. Relative mRNA levels were calculated by ΔΔCq using *TBP* expression as reference. Data were analyzed by non-parametric Wilcoxon rank-sum test.

(**F,G**) TP63 and TP73 RNA expression in tumour and CD4 T cells isolated from ATL (Acute n=22; Chronic n=24; Lymphoma n=6; Smoldering n=4), asymptomatic HTLV-1 carriers (n=3), and healthy controls (n=3). RNA-Seq data (EGAD00001001411^8^) are plotted for each donor as variance-stabilized (VST) expression, while black lines indicate median expression within each subgroup. Differential expression between each subgroup and healthy controls was tested using DESeq2.

**(H)** ChIP–qPCR enrichment over input showing p63 (α isoform) binding to the p53-responsive element^33^ within the FDXR promoter in HTLV-1^+^ cells and uninfected controls. Chromatin immunoprecipitated with anti-ΔNp63 or IgG antibodies was used as a control. Amplification products were analyzed by agarose gel electrophoresis, as shown in the lower panel. Primer sequences used to amplify the p53-responsive element are listed in Supplementary Table 4. *p <0.05, **p <0.01, ***p <0.001, ****p <0.0001.

**Supplementary Figure 5. The IRF-4/BATF3 complex binds an active enhancer region in the TP63 locus.**

**(A)** RNA-seq data from leukemia and lymphoma cell lines were retrieved from the Human Protein Atlas, stratified by FDXR expression (≥75 nTPM, Mid-High_FDXR; <75 nTPM, Mid-Low_FDXR), and used to infer a transcriptional regulatory network (regulon) as depicted in Figure 3A. Upstream transcriptional activators of TP63 (left) and TP73 (right) were extracted from this regulon. Scatter plots display, for each activator, the inferred regulatory effect on TP63 or TP73 (x-axis) versus its differential mRNA expression between Mid-High and Mid-Low FDXR groups (y-axis), estimated using limma^54^. Colours indicate the direction of differential mRNA expression of each activator (green, upregulated; red, downregulated), point size indicates regulatory strength. The top 30 transcriptional activators for TP63 and TP73 are labelled.

(**B-C**) ChIP-seq RPM-normalized signal tracks and MACS2 peak calls (q = 1e-5) for IRF4, BATF3, biotag-BATF3, H3K27ac, and BRD4 at the *TP63* locus were retrieved from published datasets in ChIP-Atlas^67^. Panel B shows the IGV visualization of TPM-normalized ChIP-seq signal tracks for ATL cell lines^35^ (ST1, KK1) and uninfected lymphoid cell lines (Jurkat, MOLT-4). MANE^76^ (v1.0; Ensembl) annotations display reference transcripts for TAp63 and ΔNp63, while FANTOM5^37^ annotations indicate experimentally defined expressed enhancer regions. Regions highlighted in light green correspond to active enhancer elements co-occupied by IRF4 and BATF3 and marked by H3K27ac and BRD4 enrichment in ATL cells. The lower zoom-in panel shows the sequence and genomic position of a canonical AICE motif (which can mediate cooperative binding of the IRF4–BATF3 complex^48^) within an active enhancer region. Genomic coordinates are based on the hg38 assembly. Panel C shows the distance between the AICE motif occurring within the TP63 locus and the summits of IRF4 and BATF3 ChIP-seq peaks in ST1 and KK1 cells. Distances were calculated using HOMER by scanning 100-bp windows centered on ChIP-seq peak summits with three previously described^48^ canonical AICE motif sequences. The sequence logo depicts the AICE motif variant identified within the TP63 locus.

**Supplementary Figure 6. Canonical T and NK lineage marker expression in integrated T/NK scRNA-seq clusters.**

Raw sequencing data from PBMCs of healthy donors and PLHTLV were retrieved from study PRJEB47382^39^. After initial clustering and lineage annotation, cells identified as T/NK lymphocytes were integrated across samples and re-clustered using PCA-based dimensionality reduction followed by Louvain clustering. The UMAP feature plots show the expression of canonical T cell (*CD3D*, *CD4*, *CD8A*) and NK cell (*GZMA*) marker genes across clusters.

**Supplementary Figure 7. Expression of marker genes used to define T/NK subset identities in scRNA-seq clusters.**

Raw sequencing data from PBMCs of healthy donors and PLHTLV were retrieved from study PRJEB47382^39^. Integrated clustering of T/NK cells was performed as shown in Supplementary Figure 6. Cluster identities were defined based on the expression of canonical lineage and functional marker genes. The dot plot displays the average expression and percentage of expressing cells for markers characterizing naïve, memory, cytotoxic, exhausted, regulatory, NK, and proliferating cell subpopulations across clusters.

**Supplementary Figure 8. Dose-dependent effects of bleomyicin and fluorouracil in vitro and ex vivo.**

(**A-C**) HTLV-1^+^/ATL and uninfected cell lines were treated with serial dilutions of bleomycin, and cell viability/proliferation was assessed by MTS assay 72 h post-treatment. Absorbance values were normalized to DMSO controls, and dose–response curves were fitted for each cell line using four-parameter logistic nonlinear regression (A,B). The fitted curves were used to calculate AUCs (C). In panel C, each dot represents a cell line and data are depicted as median ± IQR. Group-wise AUC comparisons were conducted using an unpaired Wilcoxon rank-sum test.

(**D-F**) Primary CD4 T cells from five ATL patients (1 acute, 3 chronic, 1 smoldering) and three healthy donors, along with two ATL cell lines (SP and TL-Om1), were treated with fluorouracil or bleomycin for 48 h. RNA was extracted and analyzed by Drug-Seq. Panel D shows treatment-induced pathway changes evaluated by Gene Set Enrichment Analysis (GSEA)^75^ using Hallmark gene sets, based on DESeq2 Wald statistics comparing drug-treated samples with DMSO controls. Bubble size indicates enrichment significance (−log₁₀ FDR), and color indicates normalized enrichment score (NES). The top 20 pathways (top 10 per drug by maximum |NES|) are shown for each condition. Panels E, F show lollipop plots of the top 25 genes altered by bleomycin treatment at 5 μM (E) and 10 μM (F) in primary ATL cells, based on DESeq2 analysis. Genes are ranked by statistical significance (adjusted p value). For each gene, the stem and point position indicate the log₂ fold change relative to DMSO. Point size reflects −log₁₀(adjusted p value), and point colour denotes significance (adjusted p < 0.05).

## Supplementary Tables

**Supplementary Table 1. Differential pathway analysis of proteomic data from HTLV-1**⁺**/ATL and uninfected lymphoid cell lines.** Three HTLV-1^+^/ATL (MT-2, MT-4, SP) and three uninfected (Jurkat, MOLT-4, CEM-ss) lymphoid cell lines were subjected to TMT proteomics. Each cell model was analyzed in quadruplicate. Protein expression levels were Log2 transformed and used for pathway analysis with the Camera gene set algorithm in ReactomeGSA. Pathway directionality reflects HTLV-1^+^/ATL versus uninfected comparisons.

**Supplementary Table 2. Differential expression of mitochondrial proteins in HTLV-1**⁺**/ATL and uninfected lymphoid cell lines.** Three HTLV-1^+^/ATL (MT-2, MT-4, SP) and three uninfected (Jurkat, MOLT-4, CEM-ss) lymphoid cell lines were subjected to TMT proteomics. Differential expression of mitochondrial proteins was assessed using limma and filtering the results for proteins of known mitochondrial localization (Mitocarta database^22^). Localisation reliability is derived from Human Protein Atlas annotations and classified as enhanced (highest confidence), supported, or approved.

**Supplementary Table 3. Clinical and demographic characteristics of the blood donor cohort in São Caetano do Sul, Brazil.**

**Supplementary Table 4. Sequences of primers, probes and sgRNAs used in the study.** The exons targeted by isoform specific primers/probes are shown in Supplementary Figure 4. sgRNAs were selected from the Brunello CRISPR knockout library^72^.

## References

1. Bangham, C.R.M. HTLV-1 persistence and the oncogenesis of adult T-cell leukemia/lymphoma. Blood 141, 2299–2306 (2023).

2. Valcarcel, B., et al. Prevalence and survival outcomes of adult T-cell leukemia/lymphoma in Latin America: A multicenter cohort study and recommendations to improve diagnosis and outcomes. Cancer Epidemiol 98, 102890 (2025).

3. Bangham, C.R.M., Araujo, A., Yamano, Y. & Taylor, G.P. HTLV-1-associated myelopathy/tropical spastic paraparesis (vol 1, 15012, 2015). Nat Rev Dis Primers 1(2015).

4. Cook, L.B. & Phillips, A.A. How I treat adult T-cell leukemia/lymphoma. Blood 137, 459–470 (2021).

5. Iwanaga, M., et al. Human T-cell leukemia virus type I (HTLV-1) proviral load and disease progression in asymptomatic HTLV-1 carriers: a nationwide prospective study in Japan. Blood 116, 1211–1219 (2010).

6. Saito, M., et al. Clonality analysis of HTLV-1-infected cells enhances prognostic precision in smoldering adult T-cell leukemia/ lymphoma. Haematologica 110, 2513–2517 (2025).

7. Sato, T., et al. CSF CXCL10, CXCL9, and Neopterin as Candidate Prognostic Biomarkers for HTLV-1-Associated Myelopathy/Tropical Spastic Paraparesis. Plos Neglect Trop D 7(2013).

8. Kataoka, K., et al. Integrated molecular analysis of adult T cell leukemia/lymphoma. Nat Genet 47, 1304–1315 (2015).

9. Manivannan, K., Rowan, A.G., Tanaka, Y., Taylor, G.P. & Bangham, C.R.M. CADM1/TSLC1 Identifies HTLV-1-Infected Cells and Determines Their Susceptibility to CTL-Mediated Lysis. Plos Pathog 12(2016).

10. Polakowski, N., et al. HBZ upregulates myoferlin expression to facilitate HTLV-1 infection. Plos Pathog 19, e1011202 (2023).

11. Sugata, K., et al. HTLV-1 Viral Factor HBZ Induces CCR4 to Promote T-cell Migration and Proliferation. Cancer Res 76, 5068–5079 (2016).

12. Yamagishi, M., et al. Polycomb-mediated loss of miR-31 activates NIK-dependent NF-kappaB pathway in adult T cell leukemia and other cancers. Cancer Cell 21, 121–135 (2012).

13. Sato, T., et al. Multicenter, randomized, double-blind, placebo-controlled phase 3 study of mogamulizumab with open-label extension study in a minimum number of patients with human T-cell leukemia virus type-1-associated myelopathy. J Neurol 271, 3471–3485 (2024).

14. Ishida, T., et al. Defucosylated anti-CCR4 monoclonal antibody (KW-0761) for relapsed adult T-cell leukemia-lymphoma: a multicenter phase II study. J Clin Oncol 30, 837–842 (2012).

15. Yonekura, K., et al. Mogamulizumab for adult T-cell leukemia-lymphoma: a multicenter prospective observational study. Blood Adv 4, 5133–5145 (2020).

16. Cordover, E., et al. Mogamulizumab Shows Significant Activity with Tolerable Toxicity in a Real-World North American Adult T-Cell Leukemia/Lymphoma (ATLL) Cohort. Blood 144, 6418–6419 (2024).

17. Myers, C.S., et al. Distinctive genomic features of human T-lymphotropic virus type 1-related adult T-cell leukemia-lymphoma in Western populations. Haematologica 109, 4021–4039 (2024).

18. Tanaka, A., et al. Integrative analysis of ATAC-seq and RNA-seq for cells infected by human T-cell leukemia virus type 1. Plos Comput Biol 21(2025).

19. Toyoda, K., et al. HTLV-1 bZIP Factor-Induced Reprogramming of Lactate Metabolism and Epigenetic Status Promote Leukemic Cell Expansion. Blood Cancer Discov 4, 374–393 (2023).

20. Van Weyenbergh, J., et al. Multi-cohort cross-omics analysis reveals disease mechanisms and therapeutic targets in HTLV-1-associated myelopathy, a neglected retroviral neuroinflammatory disorder. Res Sq (2025).

21. Shytaj, I.L., et al. Glycolysis downregulation is a hallmark of HIV-1 latency and sensitizes infected cells to oxidative stress. Embo Mol Med 13(2021).

22. Rath, S., et al. MitoCarta3.0: an updated mitochondrial proteome now with sub-organelle localization and pathway annotations. Nucleic Acids Res 49, D1541–D1547 (2021).

23. Slone, J.D., et al. Integrated analysis of the molecular pathogenesis of FDXR-associated disease. Cell Death Dis 11(2020).

24. Zhang, Y.H., et al. Ferredoxin reductase and p53 are necessary for lipid homeostasis and tumor suppression through the ABCA1-SREBP pathway. Oncogene 41, 1718–1726 (2022).

25. Pinto, M.T., et al. Genes related to antiviral activity, cell migration, and lysis are differentially expressed in CD4(+) T cells in human t cell leukemia virus type 1-associated myelopathy/tropical spastic paraparesis patients. AIDS Res Hum Retroviruses 30, 610–622 (2014).

26. Kodama, D., et al. Inhibition of ABL1 tyrosine kinase reduces HTLV-1 proviral loads in peripheral blood mononuclear cells from patients with HTLV-1-associated myelopathy/tropical spastic paraparesis. PLoS Negl Trop Dis 14, e0008361 (2020).

27. Araya, N., et al. Virus-induced RGMa expression drives neurodegeneration in HTLV-1-associated myelopathy. JCI Insight 10(2025).

28. Cook, L., Melamed, A., Yaguchi, H. & Bangham, C.R. The impact of HTLV-1 on the cellular genome. Curr Opin Virol 26, 125–131 (2017).

29. Alvarez, M.J., et al. Functional characterization of somatic mutations in cancer using network-based inference of protein activity. Nat Genet 48, 838–847 (2016).

30. Garcia-Alonso, L., Holland, C.H., Ibrahim, M.M., Turei, D. & Saez-Rodriguez, J. Benchmark and integration of resources for the estimation of human transcription factor activities. Genome Res 29, 1363–1375 (2019).

31. Meyer, P.E., Lafitte, F. & Bontempi, G. minet: A R/Bioconductor package for inferring large transcriptional networks using mutual information. BMC Bioinformatics 9, 461 (2008).

32. Yan, J.L., et al. BLIMP1 regulates cell growth through repression of p53 transcription. P Natl Acad Sci USA 104, 1841–1846 (2007).

33. Liu, G. & Chen, X.B. The ferredoxin reductase gene is regulated by the p53 family and sensitizes cells to oxidative stress-induced apoptosis. Oncogene 21, 7195–7204 (2002).

34. Murray-Zmijewski, F., Lane, D.P. & Bourdon, J.C. p53/p63/p73 isoforms: an orchestra of isoforms to harmonise cell differentiation and response to stress. Cell Death Differ 13, 962–972 (2006).

35. Nakagawa, M., et al. Targeting the HTLV-I-Regulated BATF3/IRF4 Transcriptional Network in Adult T Cell Leukemia/Lymphoma. Cancer Cell 34, 286–297 e210 (2018).

36. Murphy, T.L., Tussiwand, R. & Murphy, K.M. Specificity through cooperation: BATF-IRF interactions control immune-regulatory networks. Nat Rev Immunol 13, 499–509 (2013).

37. Arner, E., et al. Transcribed enhancers lead waves of coordinated transcription in transitioning mammalian cells. Science 347, 1010–1014 (2015).

38. Zhang, Y.H., et al. Ferredoxin reductase is critical for p53-dependent tumor suppression via iron regulatory protein 2. Gene Dev 31, 1243–1256 (2017).

39. Tan, B.J.Y., et al. HTLV-1 infection promotes excessive T cell activation and transformation into adult T cell leukemia/lymphoma. J Clin Invest 131(2021).

40. Hwang, P.M., et al. Ferredoxin reductase affects p53-dependent, 5-fluorouracil-induced apoptosis in colorectal cancer cells. Nat Med 7, 1111–1117 (2001).

41. Rees, M.G., et al. Correlating chemical sensitivity and basal gene expression reveals mechanism of action. Nat Chem Biol 12, 109–116 (2016).

42. Ichikawa, W., et al. Gene expression of ferredoxin reductase predicts outcome in patients with metastatic colorectal cancer treated by 5-fluorouracil plus leucovorin. Cancer Chemother Pharmacol 58, 794–801 (2006).

43. Bracht, K., Nicholls, A.M., Liu, Y. & Bodmer, W.F. 5-Fluorouracil response in a large panel of colorectal cancer cell lines is associated with mismatch repair deficiency. Brit J Cancer 103, 340–346 (2010).

44. Cooney, J.P., et al. Combination antiretroviral therapy and MCL-1 inhibition mitigate HTLV-1 infection in vivo. Cell 188, 4896–4912 e4819 (2025).

45. Ishitsuka, K., et al. Targeting Bcl-2 family proteins in adult T-cell leukemia/lymphoma: in vitro and in vivo effects of the novel Bcl-2 family inhibitor ABT-737. Cancer Lett 317, 218–225 (2012).

46. Hasegawa, H., et al. Activation of p53 by Nutlin-3a, an antagonist of MDM2, induces apoptosis and cellular senescence in adult T-cell leukemia cells. Leukemia 23, 2090–2101 (2009).

47. Li, Y., et al. p63: a crucial player in epithelial stemness regulation. Oncogene 42, 3371–3384 (2023).

48. Glasmacher, E., et al. A Genomic Regulatory Element That Directs Assembly and Function of Immune-Specific AP-1-IRF Complexes. Science 338, 975–980 (2012).

49. Chiang, M.Y. Tearing ATL apart to find HTLV’s sinister plans. Blood 135, 887–889 (2020).

50. Rauch, D.A., et al. Interferon regulatory factor 4 as a therapeutic target in adult T-cell leukemia lymphoma. Retrovirology 17(2020).

51. Takahashi, M., et al. HTLV-1 Tax oncoprotein stimulates ROS production and apoptosis in T cells by interacting with USP10. Blood 122, 715–725 (2013).

52. Shimoyama, M. Diagnostic criteria and classification of clinical subtypes of adult T-cell leukaemia-lymphoma. A report from the Lymphoma Study Group (1984-87). Br J Haematol 79, 428–437 (1991).

53. Griss, J., et al. ReactomeGSA - Efficient Multi-Omics Comparative Pathway Analysis. Mol Cell Proteomics 19, 2115–2125 (2020).

54. Ritchie, M.E., et al. limma powers differential expression analyses for RNA-sequencing and microarray studies. Nucleic Acids Res 43, e47 (2015).

55. Pang, Z., et al. MetaboAnalyst 6.0: towards a unified platform for metabolomics data processing, analysis and interpretation. Nucleic Acids Res 52, W398–W406 (2024).

56. Leek, J.T., Johnson, W.E., Parker, H.S., Jaffe, A.E. & Storey, J.D. The sva package for removing batch effects and other unwanted variation in high-throughput experiments. Bioinformatics 28, 882–883 (2012).

57. Li, H., et al. The Sequence Alignment/Map format and SAMtools. Bioinformatics 25, 2078–2079 (2009).

58. Chen, S., Zhou, Y., Chen, Y. & Gu, J. fastp: an ultra-fast all-in-one FASTQ preprocessor. Bioinformatics 34, i884–i890 (2018).

59. Dobin, A., et al. STAR: ultrafast universal RNA-seq aligner. Bioinformatics 29, 15–21 (2013).

60. Liao, Y., Smyth, G.K. & Shi, W. featureCounts: an efficient general purpose program for assigning sequence reads to genomic features. Bioinformatics 30, 923–930 (2014).

61. Love, M.I., Huber, W. & Anders, S. Moderated estimation of fold change and dispersion for RNA-seq data with DESeq2. Genome Biol 15, 550 (2014).

62. Stuart, T., et al. Comprehensive Integration of Single-Cell Data. Cell 177, 1888–1902 e1821 (2019).

63. Grandi, F., et al. popsicleR: A R Package for Pre-processing and Quality Control Analysis of Single Cell RNA-seq Data. J Mol Biol 434, 167560 (2022).

64. Germain, P.L., Lun, A., Garcia Meixide, C., Macnair, W. & Robinson, M.D. Doublet identification in single-cell sequencing data using scDblFinder. F1000Res 10, 979 (2021).

65. Aran, D., et al. Reference-based analysis of lung single-cell sequencing reveals a transitional profibrotic macrophage. Nat Immunol 20, 163–172 (2019).

66. Kogure, Y., et al. Whole-genome landscape of adult T-cell leukemia/lymphoma. Blood 139, 967–982 (2022).

67. Zou, Z., Ohta, T. & Oki, S. ChIP-Atlas 3.0: a data-mining suite to explore chromosome architecture together with large-scale regulome data. Nucleic Acids Res 52, W45–W53 (2024).

68. Robinson, J.T., et al. Integrative genomics viewer. Nat Biotechnol 29, 24–26 (2011).

69. Heinz, S., et al. Simple combinations of lineage-determining transcription factors prime cis-regulatory elements required for macrophage and B cell identities. Mol Cell 38, 576–589 (2010).

70. Katsuya, H., et al. The Nature of the HTLV-1 Provirus in Naturally Infected Individuals Analyzed by the Viral DNA-Capture-Seq Approach. Cell Rep 29, 724–735 e724 (2019).

71. Sanjana, N.E., Shalem, O. & Zhang, F. Improved vectors and genome-wide libraries for CRISPR screening. Nat Methods 11, 783–784 (2014).

72. Doench, J.G., et al. Optimized sgRNA design to maximize activity and minimize off-target effects of CRISPR-Cas9. Nat Biotechnol 34, 184–191 (2016).

73. Seashore-Ludlow, B., et al. Harnessing Connectivity in a Large-Scale Small-Molecule Sensitivity Dataset. Cancer Discov 5, 1210–1223 (2015).

74. Alpern, D., et al. BRB-seq: ultra-affordable high-throughput transcriptomics enabled by bulk RNA barcoding and sequencing. Genome Biology 20(2019).

75. Subramanian, A., et al. Gene set enrichment analysis: A knowledge-based approach for interpreting genome-wide expression profiles. P Natl Acad Sci USA 102, 15545–15550 (2005).

76. Morales, J., et al. A joint NCBI and EMBL-EBI transcript set for clinical genomics and research. Nature 604, 310–315 (2022).

