## Supplementary figures and images for "FDXR Upregulation by p63/p73 is a Prognostic and Therapeutic Marker of HTLV-1-Associated Adult T Cell Leukemia/Lymphoma"

### Supplementary Figure 1

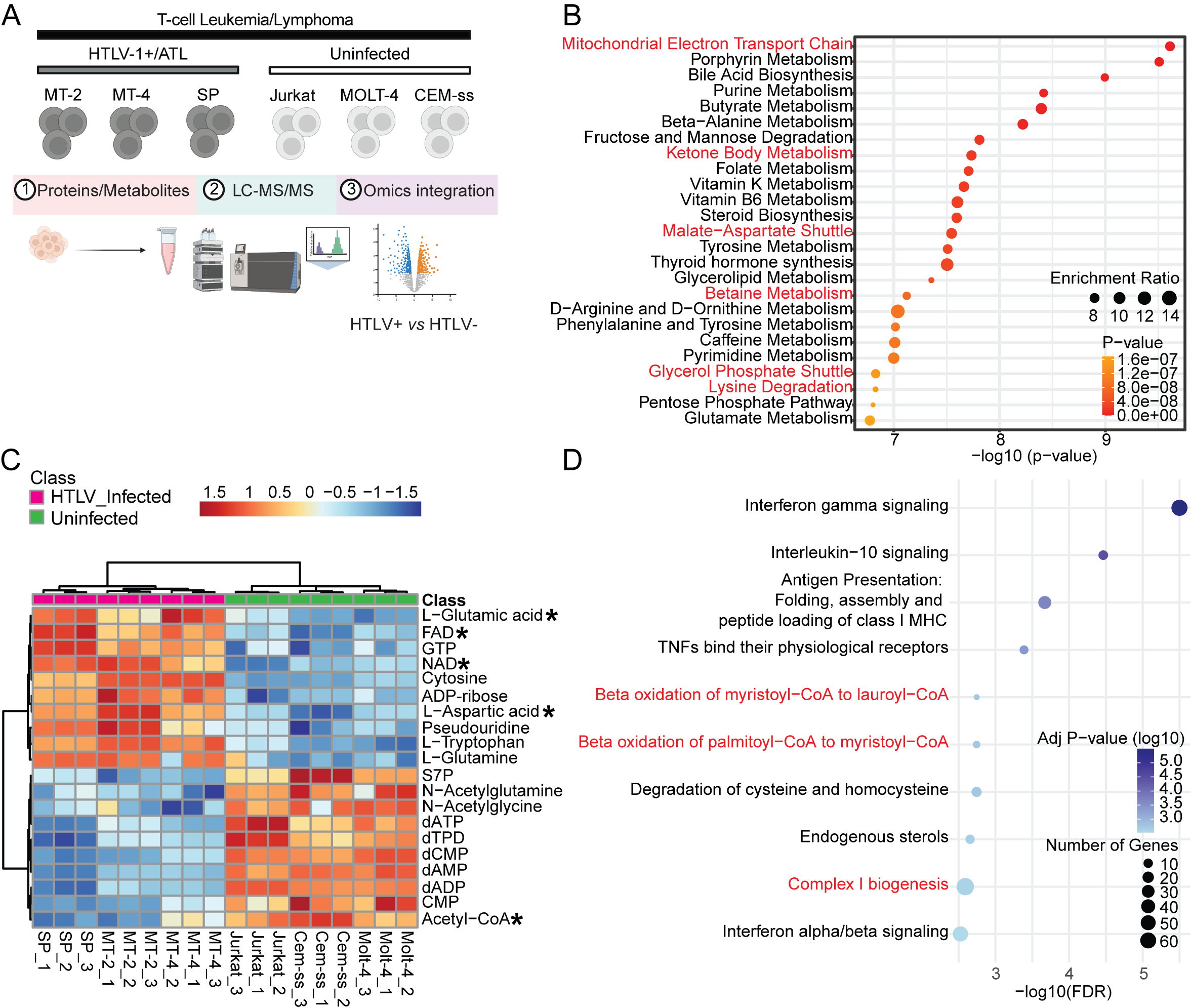

### Supplementary Figure 2

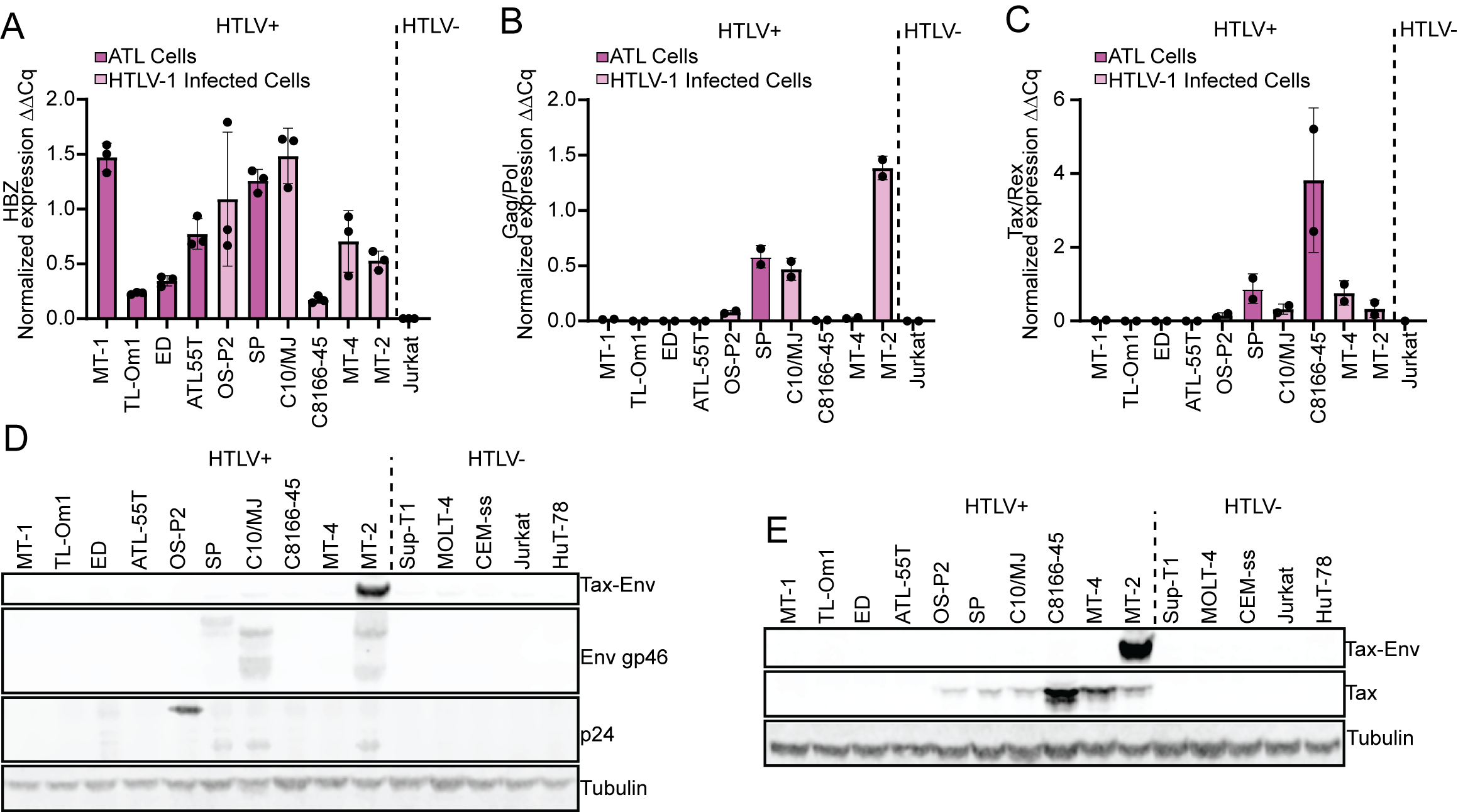

### Supplementary Figure 3

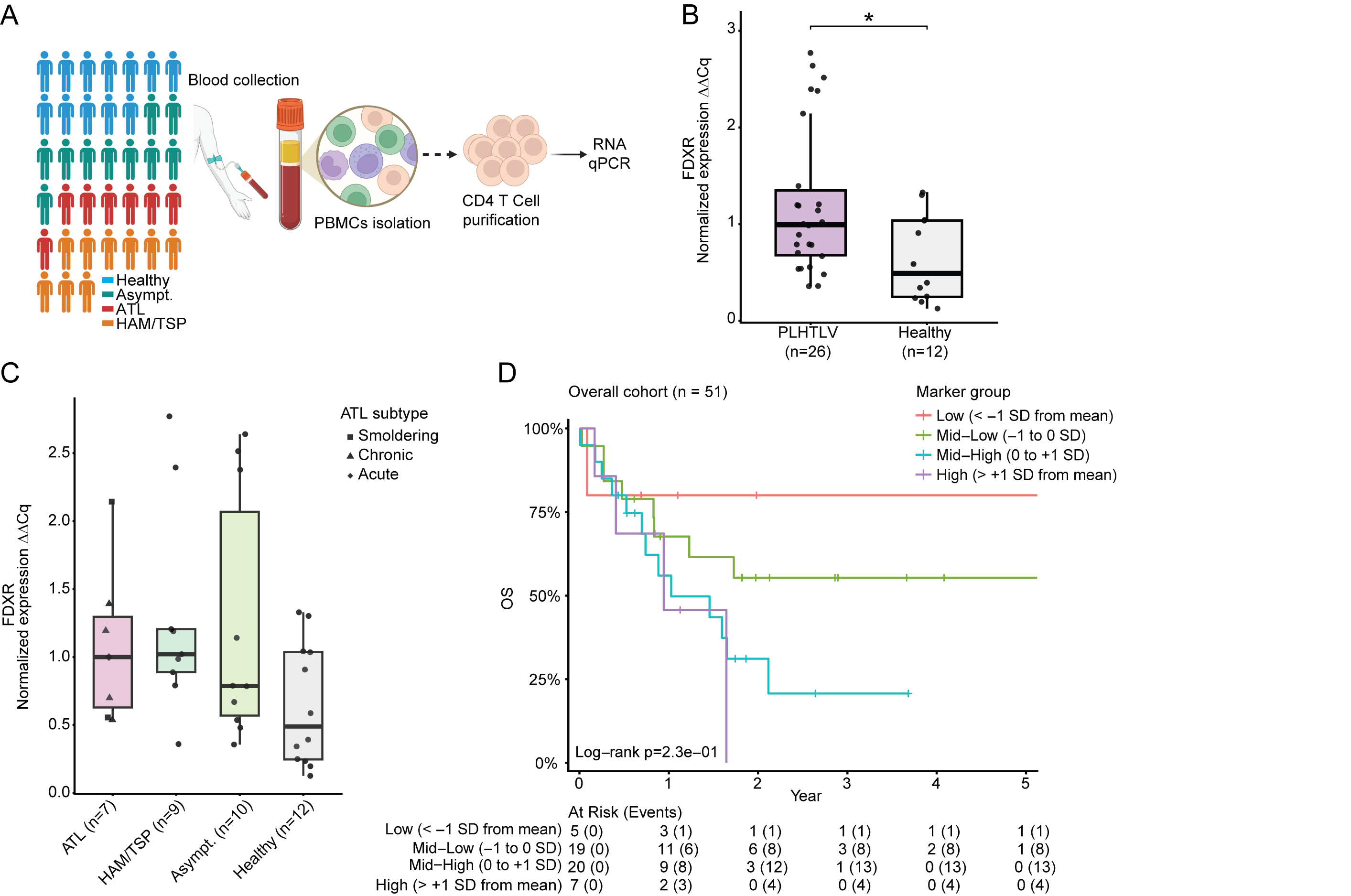

### Supplementary Figure 4

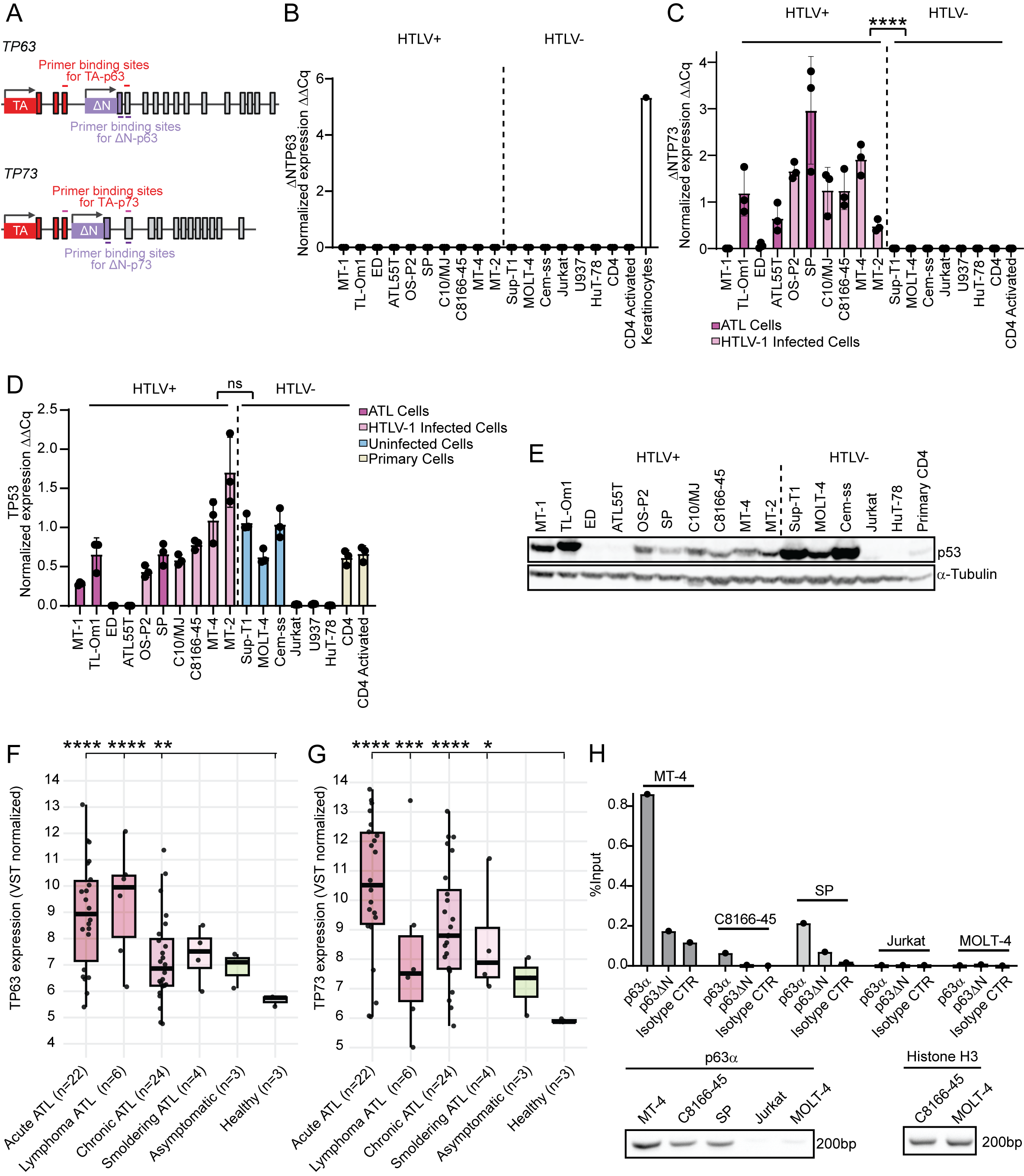

### Supplementary Figure 5

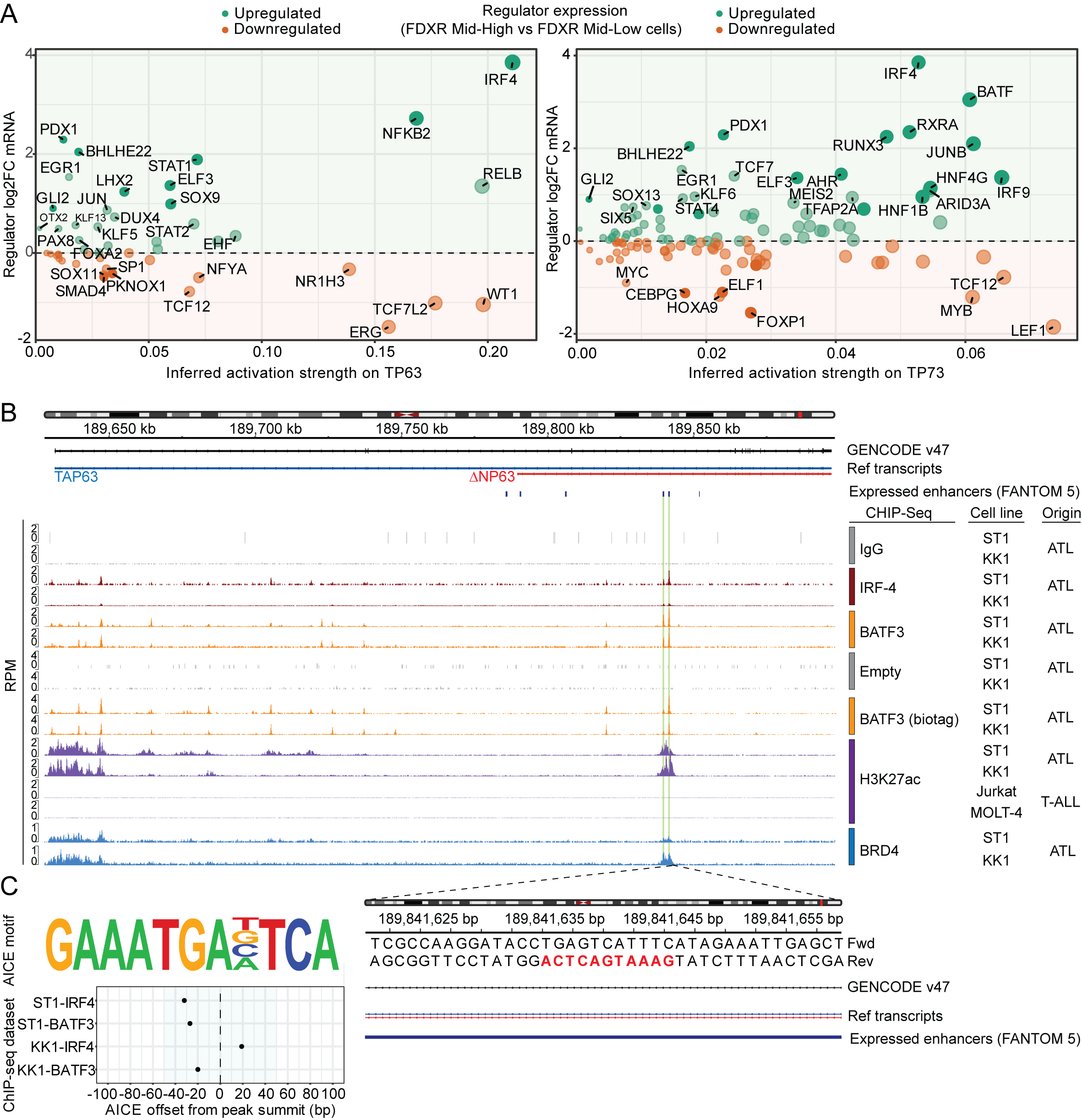

### Supplementary Figure 6

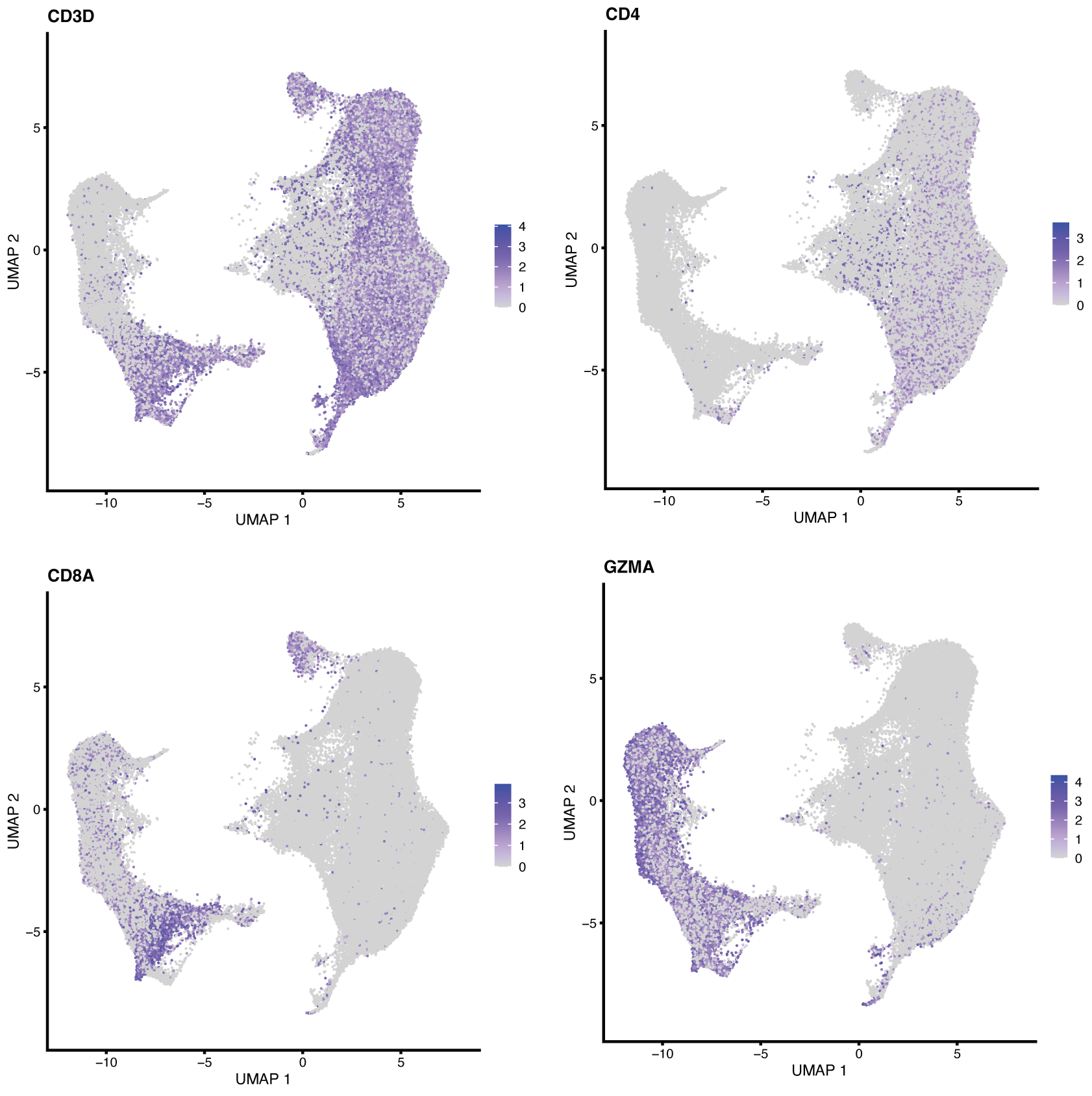

### Supplementary Figure 7

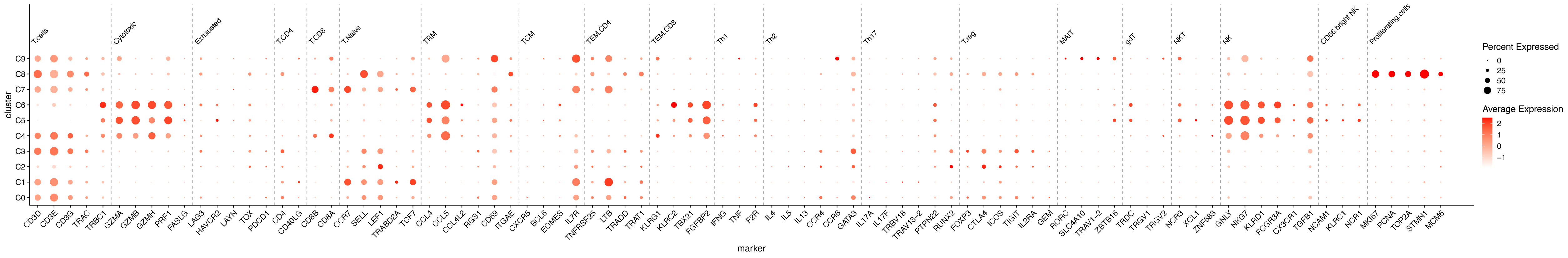

### Supplementary Figure 8

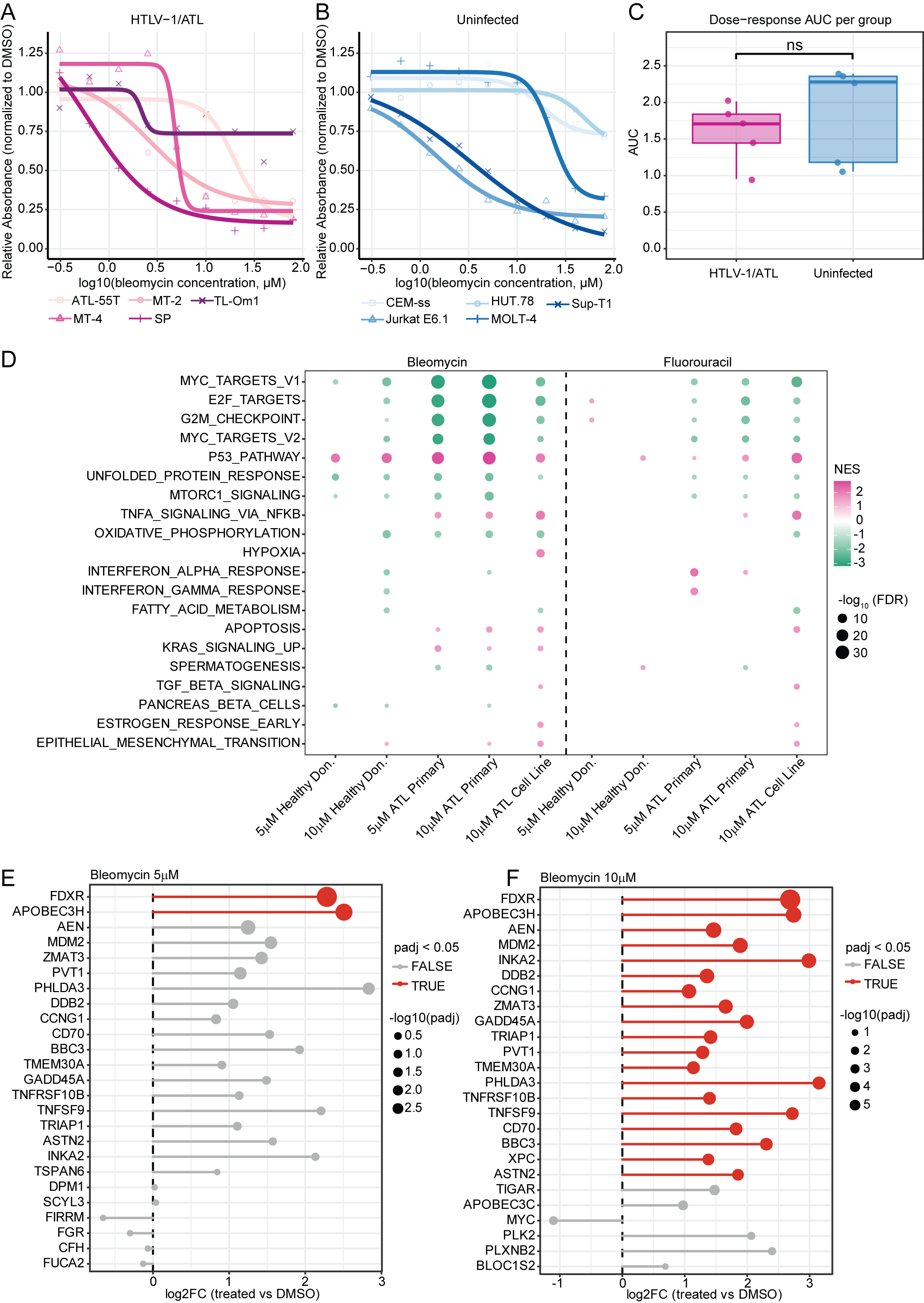
